# Resonance-organized phase-locking, extinction and phase-dependent rescue in periodically forced populations with delayed Allee effects

**DOI:** 10.64898/2026.11.23.767764

**Authors:** Junjie Sun, Frédéric Guichard, Antony R. Humphries

## Abstract

Population persistence under environmental fluctuations depends not only on the magnitude of external disturbances but also on their interaction with endogenous population dynamics. However, how environmental resonance interacts with delayed density dependence and Allee effects to determine extinction risk remains poorly understood. Here, we investigate a periodically forced Hutchinson-type population model incorporating delayed density regulation and a strong Allee effect. Through numerical simulations, Lyapunov exponent analysis, and bifurcation continuation, we characterize the organization of population dynamics and extinction across environmental forcing frequencies and amplitudes.

We find that extinction vulnerability is strongly organized by resonance between endogenous population oscillations and environmental forcing, with the lowest forcing amplitudes required for extinction occurring near the principal 1:1 resonance. The parameter space exhibits a hierarchy of frequency-locking regions consistent with Arnold tongues, including structures bounded by saddle-node bifurcations of periodic orbits. We further demonstrate that the carrying-capacity-to-Allee-threshold ratio governs the emergence of intrinsic oscillations through a Hopf bifurcation, fundamentally reorganizing the population’s response to environmental forcing. Counterintuitively, lowering the Allee threshold can increase extinction vulnerability by strengthening destabilizing delayed feedback. Finally, we identify a history-dependent rescue mechanism through which populations can recover after temporarily crossing below the nominal Allee threshold. Successful recovery events exhibit systematic phase dependence relative to environmental forcing.

Together, these findings demonstrate that extinction risk cannot be inferred from environmental severity, population abundance, or dynamical regularity alone. Instead, persistence emerges from the interaction between environmental timescales, endogenous oscillations, and history-dependent opportunities for recovery.

## 1 Introduction

Population persistence in fluctuating environments depends on both the severity of environmental variation and its interaction with population dynamics. Periodic changes in resource availability can reinforce, suppress or synchronize oscillations generated by density-dependent regulation. Consequently, populations exposed to environmental fluctuations with comparable means and amplitudes may respond differently when the period of environmental forcing changes relative to their endogenous timescales. Such interactions have long been recognized in delayed population models with periodically varying carrying capacity(Nisbet and Gurney 1976) For populations exhibiting cyclic dynamics, the timing of environmental change can also influence whether a disturbance leads to persistence or collapse(Alkhayuon et al. 2021). These observations motivate an examination of how environmental periodicity organizes extinction risk, beyond the effects of mean environmental conditions alone.

The ecological significance of these external-endogenous interactions depends on how the resulted population fluctuations interact with mechanisms of extinction, a common such case being density dependent extinctions. The Allee effect describes positive density dependence at low population densities, where individual fitness or per-capita population growth declines as populations become small (Courchamp et al. 1999; Stephens et al. 1999). Under a strong Allee effect, population growth becomes negative below a critical density threshold, creating an unstable equilibrium below which populations tend toward extinction. Oscillations can therefore threaten persistence by bringing a population into a low-density region from which recovery is difficult. Previous theoretical work has demonstrated that Allee effects can interact with complex population dynamics to produce extinction and prolonged transient behaviour(Schreiber 2003).

Delayed density dependence provides one way through which populations develop endogenous oscillations that can resonate with external environmental conditions. In Hutchinson-type models, population regulation depends on past abundance, representing a lag between population density and its consequences for subsequent growth, Ecologically, the delay represents processes such as maturation, reproduction, delayed resource feedback. The Hutchinson delayed logistic model extends logistic growth by allowing density dependence to act through a delay term in its component and has become a canonical delay differential equation in population dynamics (Hutchinson 1948) that is rigorously analyzed and studied. Its equivalent Wright formulation motivated foundational analyses of stability, oscillation, and Hopf bifurcation, with the positive equilibrium losing stability at the critical threshold(Wright 1955), resulting in a characteristic oscillation. Classical blowfly experiments by Nicholson provided an important empirical example of pronounced delayed population oscillations(Nicholson 1954; Gurney et al. 1980) Later work resolved the long-standing Wright global-stability conjecture(van den Berg and Jaquette 2018). In delayed Allee models, persistence can depend on the preceding population history, and the nominal Allee equilibrium need not act as an instantaneous threshold that determines the outcome of every trajectory(Liz et al. 2015). Understanding extinction under periodic forcing therefore requires attention to both low-density excursions and the histories that permit or prevent subsequent recovery.

An important type of interaction that can be generated between external forcing and endogenous cycle is resonance induced synchronization. Under sufficiently strong forcing, the internal cycles created by delay component act as an oscillator and may become frequency- and phase-locked to the driver, including at rational p:q frequency ratios; the corresponding locking regions in forcing-frequency–amplitude space form characteristic Arnold tongues(Pikovsky et al. 2001). Such structures provide a general framework for distinguishing synchronized periodic dynamics from quasiperiodic responses.Resonance tongues have been well demonstrated in delay differential equations: in delayed ENSO models, interactions between delay-generated intrinsic oscillations and seasonal forcing produce multiple locking regions, tongue overlap, and transitions to complex or chaotic dynamics (Keane et al. 2015; Keane et al. 2016). More broadly, Arnold-tongue analysis has been applied across climate dynamics, neuroscience, biological clocks, and optical and mechanical oscillator systems.

Despite being a ubiquitous phenomenon in nonlinear dynamical systems, resonance has received comparatively scattered attention in ecological modelling, with studies distributed across particular population, predator–prey, community, and host–pathogen systems rather than forming a unified ecological framework. Periodically forced predator–prey models can exhibit frequency locking, quasiperiodicity, chaos, and well-defined Arnold tongues, demonstrating that intrinsic population cycles may synchronize with environmental forcing at multiple frequency ratios (Allen 1990; Vandermeer et al. 2001). Experimental work further showed that periodic resource fluctuations can resonate with intrinsic population dynamics and substantially alter mean abundance in protist populations (Orland and Lawler 2004). Phase locking has additionally been demonstrated experimentally in spatial predator–prey communities, where dispersal synchronized otherwise drifting population cycles (Vasseur and Fox 2009). More recently, environmental frequency locking was shown to suppress large oscillations and promote persistence in theoretical phage– bacteria systems(Luo et al. 2026), highlighting the potential importance of resonance for ecological stability and extinction risk.

Existing studies provide complementary perspectives on interactions between forcing, delay and Allee effect. Analyses of delayed Allee models have established effects on equilibrium stability, oscillations and extinction basins(Jankovic and Petrovskii 2014; Liz et al. 2015), while periodically varying carrying capacity has been investigated in discrete population models with Allee effects(Chugunova and Ann Campbell 2020). Research on forced delay equations provides a framework for identifying the resonance structures that organize oscillatory responses(Keane et al. 2015). However, the existence of frequency locking does not, by itself, determine its consequences for persistence. In a phage–bacteria model, environmental locking can suppress oscillatory extinction(Luo et al. 2026), illustrating that synchronization may protect populations under some extinction mechanisms. This raises a corresponding question for populations with strong Allee effects: how do resonance structures organize density-dependent extinction boundaries, and consequently determines population persistence?

Here, we investigate these questions using a single-species delay differential equation that combines Hutchinson-type density regulation, a strong Allee effect and periodic variation in carrying capacity. Both density-dependent factors depend on past population abundance, whereas carrying capacity varies with the current environmental phase. We examine how do resonance regions organize the boundary between persistence and extinction as forcing amplitude and frequency vary across the parameter space, and look into the mechanisms through which extinction boundary is determined and how population behaves around it.We hope to provide an explanation the relation between this geometry of resonance and extinction-rescue dynamics.

## 2 Model formulation

### 2.1 Delayed Allee–logistic equation

Population abundance N(t) is governed by

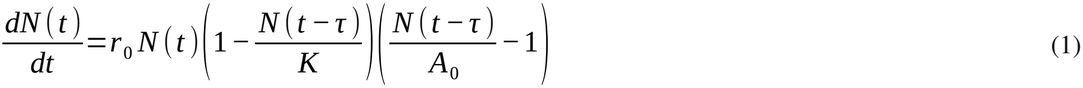

where r0 is the baseline intrinsic rate, *τ* is a fixed delay, A0 is the strong Allee threshold and K(t) is the environmental carrying capacity. Only the density-dependent feedback is delayed; the multiplicative abundance N(t) is current. The default history is the constant function Nh(s) = N0 on −*τ* ≤ s ≤ 0.

The carrying capacity is forced sinusoidally as

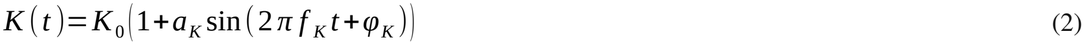

Here K0 is the mean carrying capacity, aK is a dimensionless relative amplitude, fK is ordinary frequency in cycles per model-time unit, φK is phase and ωK = 2πfK is angular frequency. The absolute forcing amplitude is BK = K0aK, which keeps every term in K(t) in population units.

### 2.2 Equilibria and the first Hopf threshold

When aK = 0, the three constant equilibria are N* = 0, A0 and K0. For K0 > A0, extinction is locally stable and the Allee equilibrium is unstable. Linearization about K0 gives

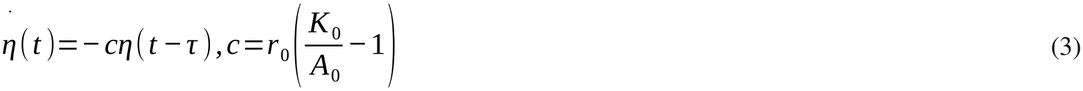

with characteristic equation μ + c exp(−μ*τ*) = 0. The first imaginary-root crossing occurs at

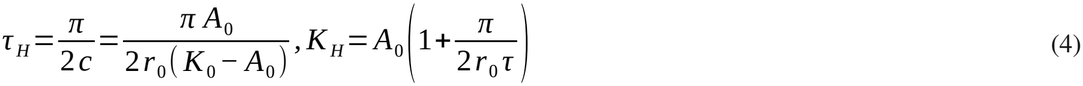

At r0 = 1, K0 = 500 and A0 = 100, *τ*H = π/8 ≈ 0.3927. The baseline *τ* = 0.5 is therefore beyond the first loss of equilibrium stability. Equivalently, at *τ* = 0.5 the frozen carrying-capacity threshold is KH ≈ 414.16. Under forcing, K(t) changes from an equilibirum to a periodic function, the meaning of the hopf threshold preserves only when N(t) is tracking K relatively closely and K(t) varies slow enough.However we find the parameter meaningfull in predicting parameter space behaviour in result part even when such condition is not met.

### 2.3 Nondimensionalized model

To identify the independent parameter combinations governing the system’s dynamics and facilitate comparisons across different carrying capacities and Allee thresholds, we introduce the dimensionless population abundance and time variables

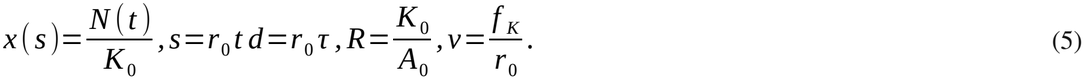

Substituting these variables into Eq. (1) yields the nondimensionalized delayed Allee–logistic model

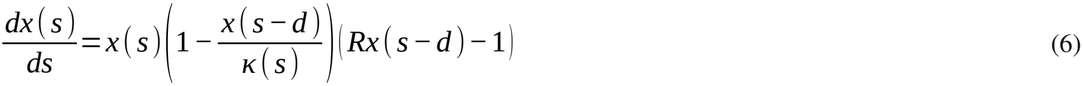

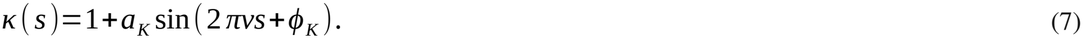

The dimensionless initial history is given by (*x*_*h*_ (*s*)=*N*_*h*_ (*s* / *r*_0_)/ *K* _0_) *for* (*s* ∈[−*d*, 0]). Under this transformation, the mean carrying capacity is normalized to unity, while the Allee threshold becomes (*x* _*A*_=1/ *R*).

For the unforced system, linearization around the positive equilibrium (x^*=1) gives

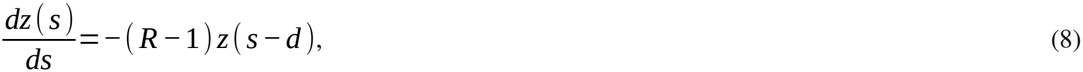

so that the strength of delayed feedback is characterized by the combined parameter (d(R-1)). The first Hopf bifurcation occurs at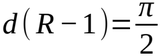.This provides an analytical reference for examining how the carrying-capacity-to-Allee-threshold ratio modifies intrinsic oscillatory dynamics and their response to periodic environmental forcing.

## 3 Numerical and figure-generation methods

### 3.1 Baseline simulation design

This is the default parameter to all constructed heatmaps unless otherwise mentioned. The empirically observed frequency f0 is measured from an unforced post-transient trajectory.We choose a parameter space range that would best demonstrate the interaction of the Forcing, delay and Allee model based on preliminary trials. (Table 1).

**Table 1.** Baseline parameters and numerical controls for amplitude–frequency banks. Values apply unless otherwise specified.

| **Quantity** | **Baseline** | **Role in the implemented banks** |
|---|---|---|
| r0 | 1.0 | Intrinsic rate |
| K0 | 500 | Mean carrying capacity |
| A0 | 100 | Allee threshold |
| τ | 0.5 | 50 numerical steps |
| Nh(s) | 300 | Constant history on [−τ, 0] |
| Δt | 0.01 | Fixed Heun step |
| Ttotal | 500 | 50,000 steps for the displayed banks |
| Ttransient | 200 | First 20,000 steps discarded |
| Next | 10<sup>−6</sup> | Stored extinction threshold |

### 3.2 Fixed-step delay integration

The model uses an improved Euler (Heun) predictor–corrector. Let Fi = F(Ni, Ni−d, ti), where d = *τ*/Δt = 50. One step is

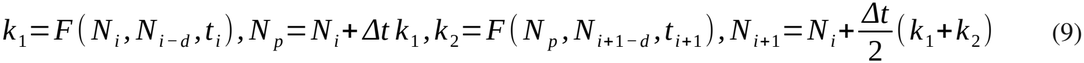

If a requested delayed index precedes t = 0, the constant history Nh = 300 is used. This makes the physical delay exactly dΔt. Parameter values and trajectories are stored in HDF5 groups so that classification and plotting do not require reintegration.

### 3.3 Construction of dynamics maps and periodicity detection

Each colored cell in corresponds to one simulated pair (aK, fK) and one such run. After the transient, the workflow checks extinction and then tests whether the delayed state profile repeats after q forcing periods, q = 1,…,8. In the implemented history-state comparison.In the most figures presented in this paper, parameter space run behaviour is classified into periodic(q=1-8), quasiperiodic, chaostic and extinction.Examples of runs classified in these cases are demonstrated in figure 10.1-4 (Appendix C.1), also see periodicity detection in the appendix. Detailed operational definitions are given in Appendix B.1–B.2 (Eqs. (17)–(22)).

**Fig. 1.**
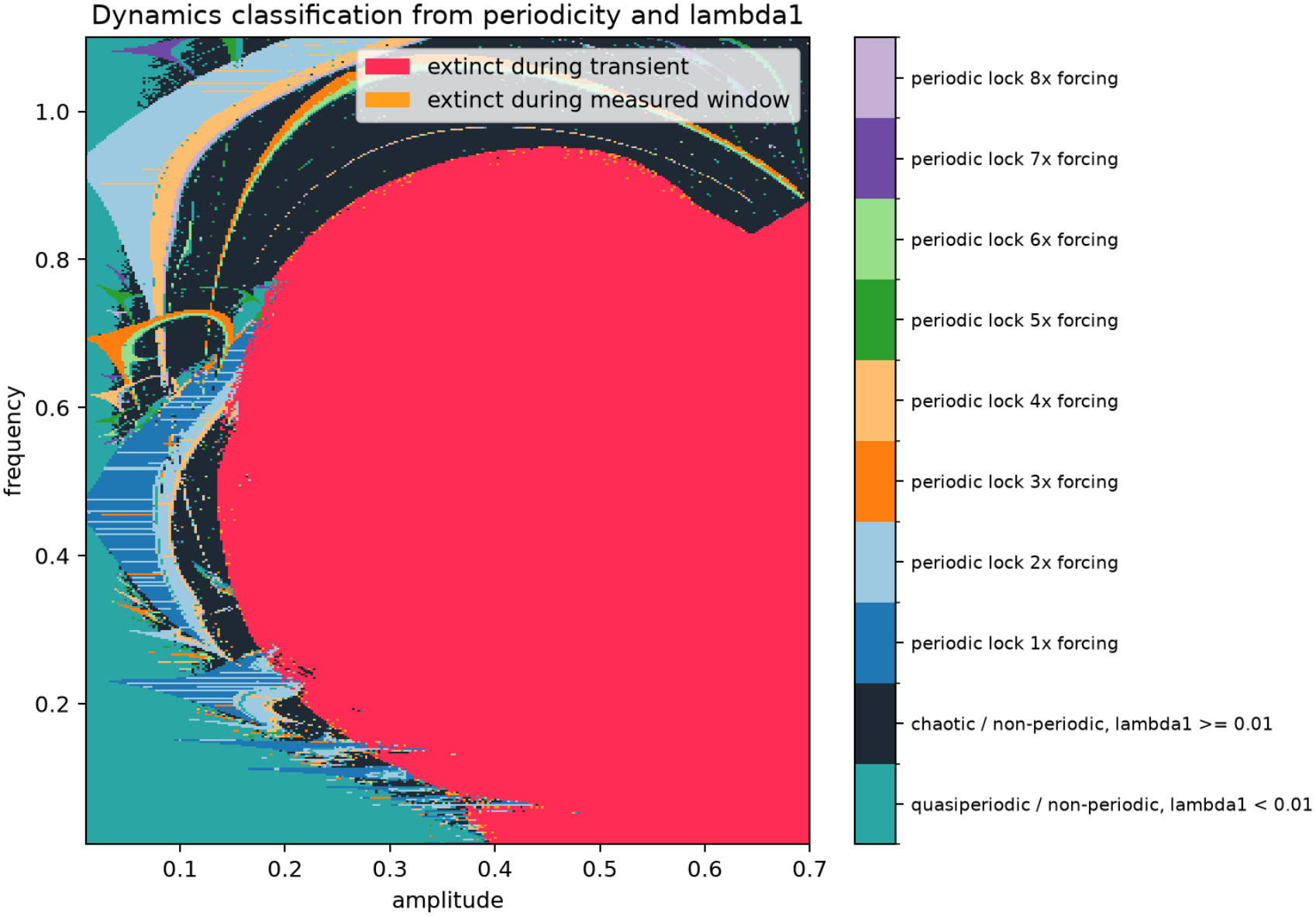
Amplitude–Frequency Parameter space Run classification map. Each cell is one fixed-history integration of Eqs. (1)–(2) at the displayed relative carrying-capacity amplitude aK and ordinary frequency fK. After 200 model-time units are discarded, the smallest accepted repeat q = 1,…,8 gives the period-q color. Cyan marks a low-Lyapunov nonperiodic candidate and dark grey a positive-Lyapunov candidate using the λ1 = 0.01 threshold printed in the stored legend. Pink and orange denote extinction before and after the transient cutoff. The colored bands are finite-grid locking candidates; the extinction wedge records the outcome from Nh = 300 and is not a continued bifurcation boundary.

### 3.4 Lyapunov exponents and Floquet multipliers

We estimate the two leading Lyapunov exponents by integrating analytic variational equations along a reference trajectory and periodically applying QR orthogonalization to perturbation histories. This follows the tangent-vector orthogonalization framework of Benettin and colleagues. The history discretization, integration scheme and numerical settings are specific to this implementation.Schematically, after repeated perturbation growth and renormalization, (Benettin et al. 1980). (Benettin et al. 1980; Breda and Liessi 2024).

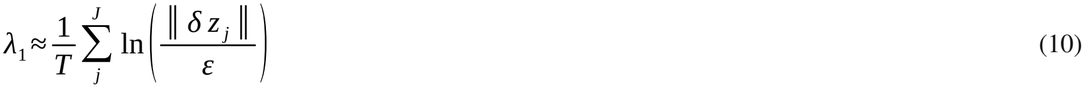

with ε the reset perturbation norm. A run with *λ*_1_>0.01 is classified as chaostic, otherwise, it is classified as quasiperiodic if not periodic. See a detailed implementation in appendix The complete tangent-history scheme is provided in Appendix B.3; the discrete Floquet implementation in Appendix B.4.

### 3.5 Log-growth mechanism and phase statistics

To investigate the mechanism underlying population rescue following severe declines, we examine how delayed density dependence and periodic environmental forcing jointly determine population growth. In particular, we seek to distinguish temporary population decline from sustained negative growth leading to extinction, and to identify the conditions under which a declining population can subsequently recover.

We define that such an event as a successful rescue event: when population density K(t) drop below Allee threshold A, but was able to recover back above A at a later time point.

Since population persistence depends on the cumulative effects of positive and negative growth over time, we express the population dynamics in terms of the instantaneous per-capita growth rate.

For N(t) > 0, Eq. (1) can be written as d ln N/dt = q(t), where

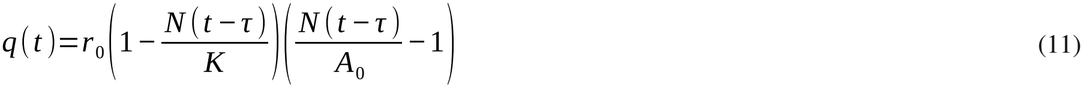

The sign rule is exact: q(t) > 0 if and only if A0 < N(t − *τ*) < K(t); q(t) < 0 when N(t − *τ*) < A0 or N(t − *τ*) > K(t). However, instantaneous negative growth does not necessarily imply extinction. Integrating the per-capita growth rate over an interval [ta, tb],

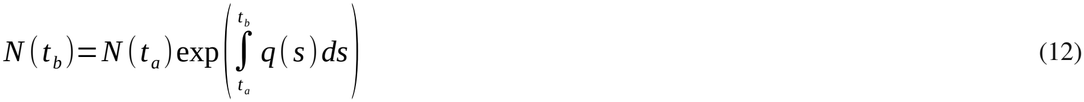

This relationship shows that changes in population abundance depend on interaction of accumulated log growth and current environmental condition, rather than the instantaneous growth rate at any single time. At the successful rescue event starting at a and ending at b, *N* (*t*_*b*_)=*N* (*t*_*a*_).

These quantities allow us to examine whether successful rescue is associated with sufficient compensatory growth following environmental deterioration.

We further investigate whether successful rescue depends on the timing of the disturbance within the environmental forcing cycle. Because the carrying capacity varies periodically, the timing of a disturbance relative to the forcing cycle may influence subsequent population recovery. To characterize this phase dependence, we record the carrying-capacity phase at the downward crossing associated with each successfully rescued event (j)

For each successfully rescued event j, the carrying-capacity phase at the downward crossing is

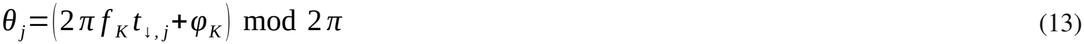

We then calculate the circular mean direction and circular variance of the phases of successful rescue events:

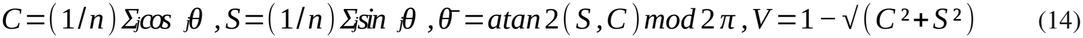

Here, (n) denotes the number of successful rescue events within a simulation. The circular mean direction *θ* _*j*_ characterizes the typical environmental phase associated with successful rescue, while the circular variance (V) quantifies the degree of phase concentration. Values of (V) approaching zero indicate that rescue events occur at similar phases, whereas values approaching one indicate weak phase concentration.

Together, these measurements characterize two complementary aspects of the rescue process: the balance of accumulated population growth during recovery and the temporal organization of successful rescues relative to periodic environmental forcing. The circular mean direction are presented in Figures 7

## 4 Results

### 4.1 Environmental frequency and forcing amplitude organizes extinction through resonance induced tongue patterns

#### 4.1.1 Resoanance organized extinction boundary

We first characterized population responses to periodic carrying-capacity forcing across the amplitude– frequency parameter space using stroboscopic periodicity detection and the largest finite-time Lyapunov exponent (Figure 1). The forcing frequency strongly influences the observed extinction outcomes. The broad early-extinction domain approaches smaller forcing amplitudes in the neighborhood of the principal 1:1 resonance (Figures 1 and 2a), where external forcing frequency equals intrinsic oscillation frequency of the unforced population, measured as (f_0_≈0.455). Above and below this region, increasing the difference between intrinsic and forcing frequencies generally shifts the early-extinction boundary toward larger forcing amplitudes.

**Fig. 2.**
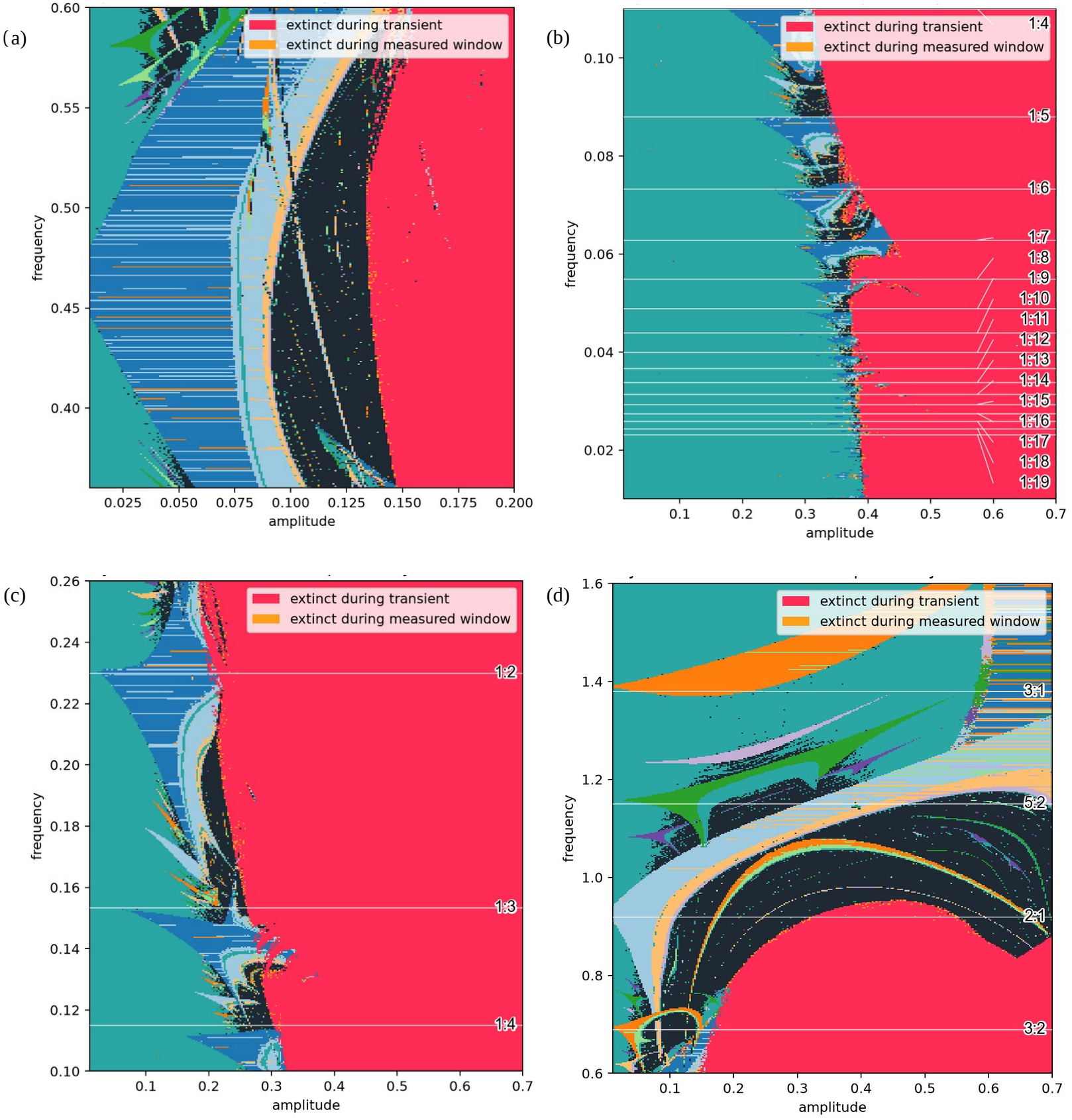
Largest Tongue steming from pinciple resonance The four panels resolve distinct frequency ranges: (a) the principal near-1:1 response, (b–c) lower-frequency locking structures, and (d) higher-frequency responses. The colored cells retain the classification and extinction coding of Fig. 1. Panel-specific frequency limits are shown on their axes.

A pronounced asymmetry is observed in extinction boundary across the principal resonance frequency. At higher forcing frequencies,extinction boundary retreats much sharper, leaving the parameter space into prolonged persistence accompanied by chaostic dynamics. At lower frequencies, extinction boundary is surrounded by bits of locking and chaostic regions intersecting each other, also accompanied by a smoother retreat of extinction boundary into higher amplitude regions.These results highlights the role of principal resonance as a boundary in different parameter space behaviour and pivot in extinction organization.

#### 4.1.2 Arnold’s togues in parameter space

Under weak forcing, much of the parameter space is occupied by low-Lyapunov nonperiodic responses, interspersed with wedge-shaped regions of periodic locking. These regions generally broaden over a greater range of forcing frequencies as forcing amplitude increases, forming a hierarchical pattern of frequency-dependent dynamical responses. Positive-Lyapunov(Chaostic) regions frequently appear around the boundaries of the locking regions and where multiple structures approach one another. At sufficiently large forcing amplitudes, extensive portions of parameter space transistions to extinctions.

The most prominent locking region originates near the intrinsic oscillation frequency of the unforced population, measured as (*f* _0_ ≈ 0.455) (Figure 2a). This region broadens rapidly with increasing forcing amplitude and is consistent with the principal 1:1 resonance between intrinsic population oscillations and external environmental forcing. Additional locking regions occur near rational frequency ratios relative to (*f* _0_), including lower forcing-frequency ratios of approximately 1:2, 1:3, and 1:4, together with higher-frequency ratios such as 2:1 and 3:1 (Figure 2b–d). Finer inspection of the low-frequency region reveals a sequence of increasingly narrow structures associated with higher-order rational ratios.

Further investigation of the 2:1 phase-locking tongue (Figure 3) confirms that the periodic locking region is bounded by fold (saddle-node) bifurcation curves of periodic orbits. Along these boundaries, a Floquet multiplier crosses the unit circle at +1, indicating a change in the stability of the corresponding periodic solutions. This is consistent with the classical saddle-node mechanism underlying the formation of Arnold tongues, providing bifurcation-based confirmation of the observed phase-locking structure.

**Figure 3.**
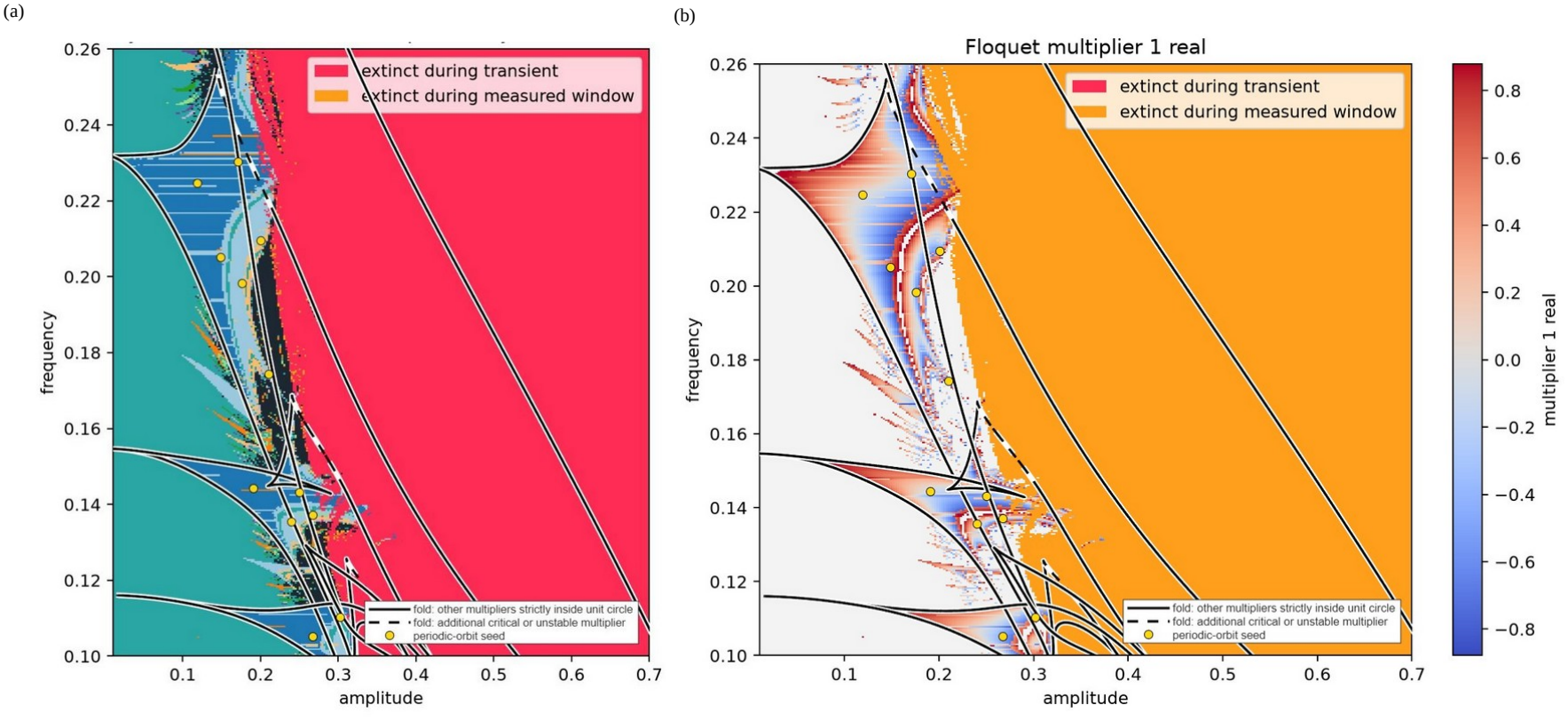
Numerically continued fold (saddle-node) bifurcation curves of periodic orbits superimposed on (left) the dynamical regime classification map and (right) the real part of the first Floquet multiplier. Solid curves indicate fold bifurcations for which all remaining Floquet multipliers lie strictly within the unit circle, whereas dashed curves indicate the presence of additional critical or unstable multipliers. Yellow markers denote periodic-orbit seeds used for numerical continuation.

Overall, these observations indicate that intrinsic–external frequency relationships provide an important organizing framework for the forced dynamics. Resonance-associated structures are closely related to the distribution of periodic locking, irregular population oscillations, and extinction, motivating further investigation of their bifurcation mechanisms and ecological consequences.

### 4.2 The Allee threshold, Hopf bifurcation threshold, and their relation to extinction booundary

To investigate how the Allee effect influences resonance organization and extinction boundary, and to determine whether it affects such determination on its own or through an intergrative component, we produces heatmaps with varying Allee threshold.

Linear stability analysis gives the Hopf condition

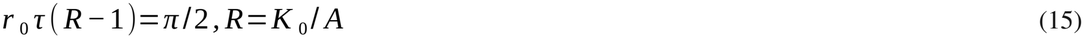

which yields a critical ratio of (*R*_*h*_ ≈ 4.14) for (*r* _0_=1) and (*τ* =0.5). We first fixed (*K* _0_=500) and progressively decreased (A), corresponding to an increase in (R) from 3.00 to 5.50 (F01–F07; Fig. 4) around the hopf bifurcation thershold (between F03 and F04). Then we identify two functional component,(R) value, and hopf-bifurcation component *r* _0_ *τ* (*R*− 1) that includes (A), and reverse parts other than A in these components to see how much of the dyanmical region behaviour is returned to baseline.

**Fig. 4.**
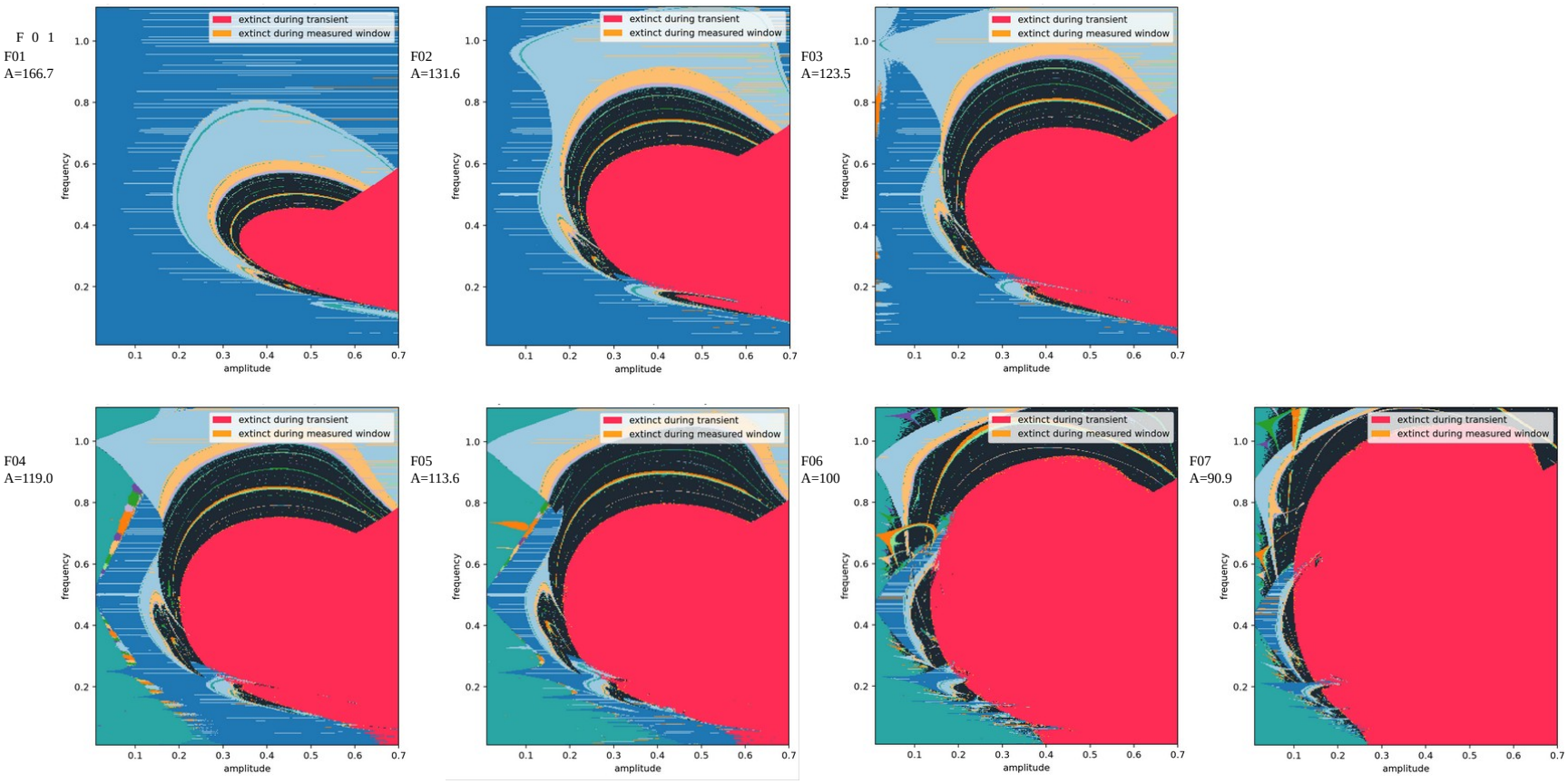
F-series response maps at fixed K_0_ = 500, r_0_ = 1, and N_0_ = 300 while the Allee threshold decreases from A = 166.7 (F01) to 90.9 (F07), corresponding to R = K_0_/A increasing from 3.00 to 5.50. The seven panels show the appearance of distinct locking bands across the unforced first-Hopf threshold (R ≈ 4.14). Colors follow the dynamics classification in Fig. 1.

For (R) component, K02, K04 and K07(Fig. 5) has same (A) value as F02,F04 and F07, yet identical (R) component value as base line F06 by varying(K). For hopf bifurcation component, M015, M020,M030(Fig. 6) has different (A) value with each other and same *r* _0_ *τ* (*R*− 1) by varying (r)

**Fig. 5.**
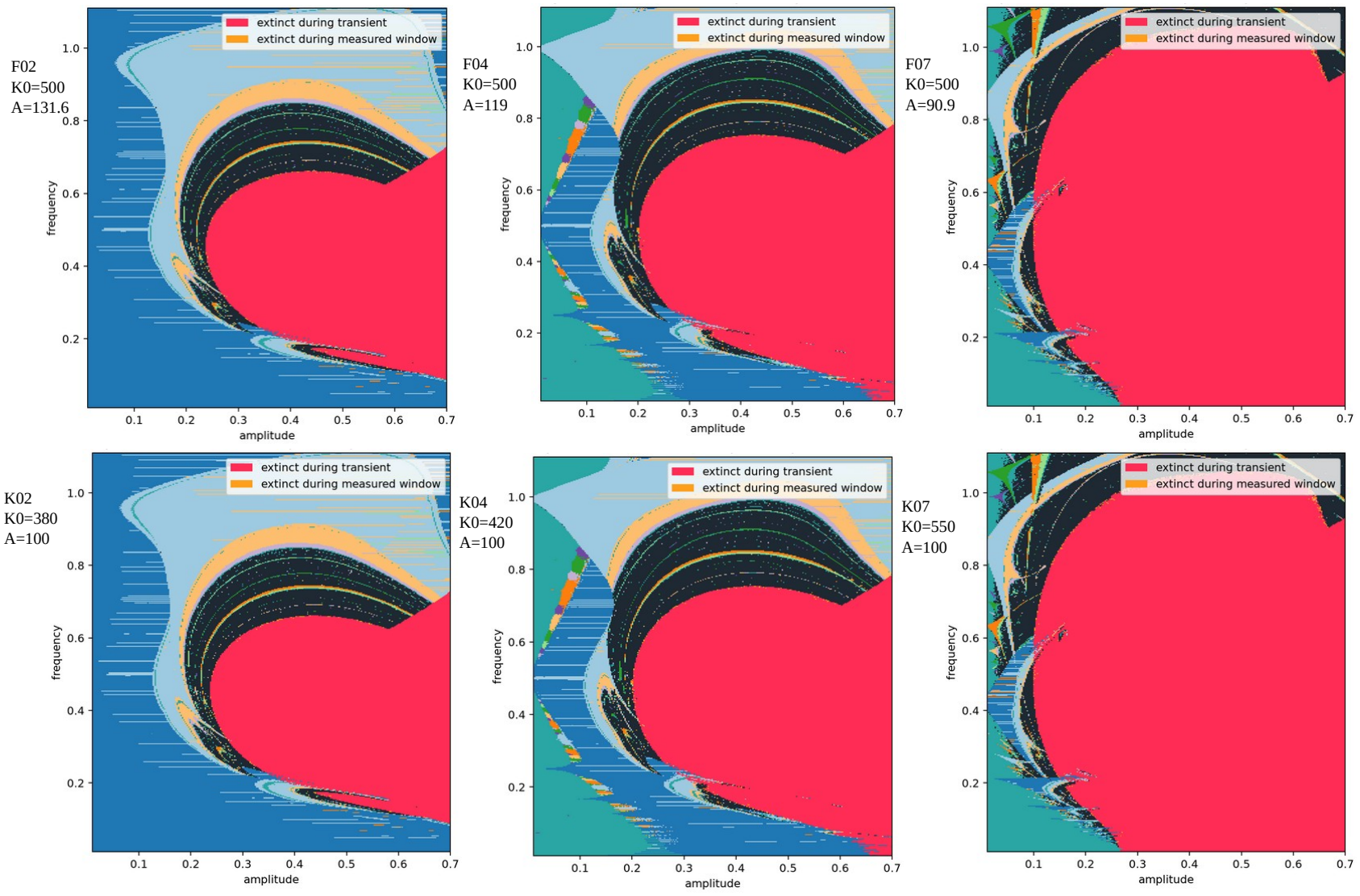
Carrying-capacity compensation experiment with paired F- and K-series maps at matched ratios R = 3.80, 4.20, and 5.50. The first row retains K_0_ = 500 and varies A; the second row holds A = 100 and varies K_0_ with scaled initial abundances. The comparison assesses invariance under changes in the absolute abundance scale. Colors follow Fig. 1.

**Fig. 6.**
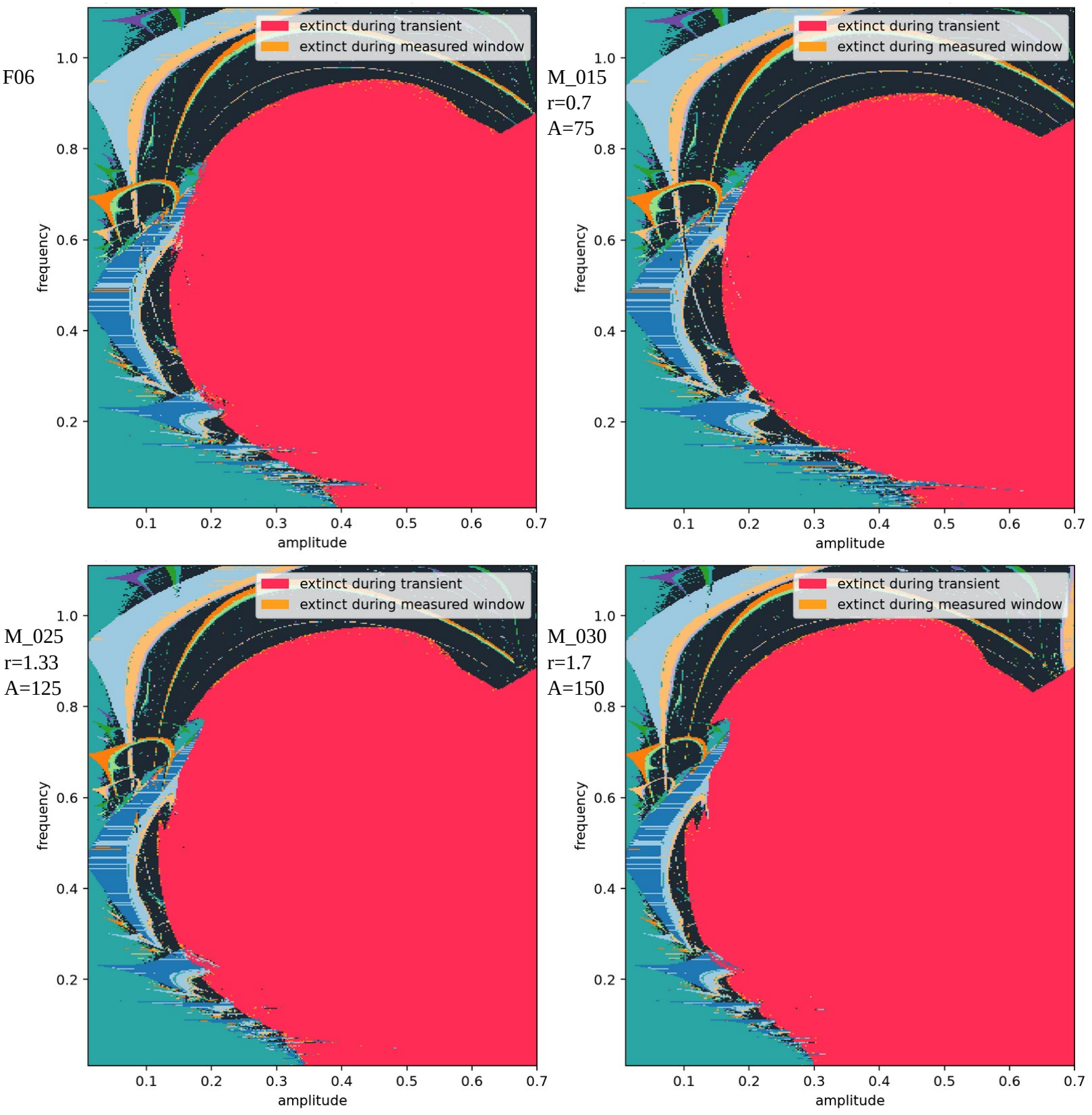
Compensatory growth-rate and Allee-threshold experiment. The reference bank F06 (r_0_ = 1, A = 100) is compared with M_015 (r_0_ ≈ 0.706, A = 75), M_025 (r_0_ ≈ 1.333, A = 125), and M_030 (r_0_ ≈ 1.714, A = 150) at K_0_ = 500 and N_0_ = 300. Each combination approximately preserves c = r_0_(K_0_/A − 1). Colors follow Fig. 1.

**Fig. 7.**
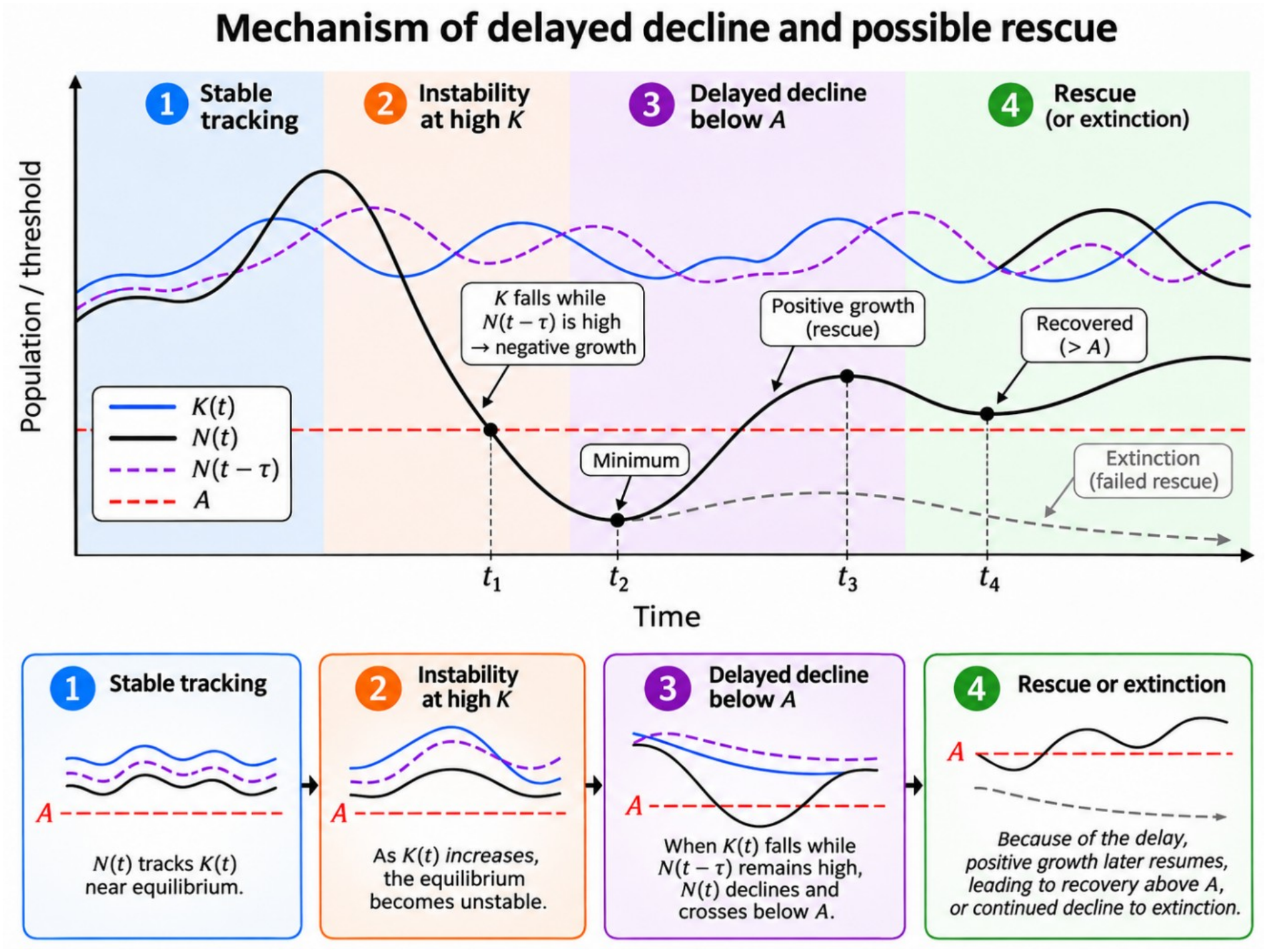
Mechanistic schematic of delayed decline and possible rescue. Panel A shows an illustrative current population N(t) (black), carrying capacity K(t) (blue), delayed population N(t − *τ*) (purple dashed), Allee threshold A0 (red dash-dot) and a failed-rescue branch (grey dashed). Growth is positive only when A0 < N(t − *τ*) < K(t). The labels t1–t4 summarize a downward threshold crossing, a minimum, a later maximum and a secondary minimum. Panel B condenses this sequence into conceptual stages.

**Fig. 8.**
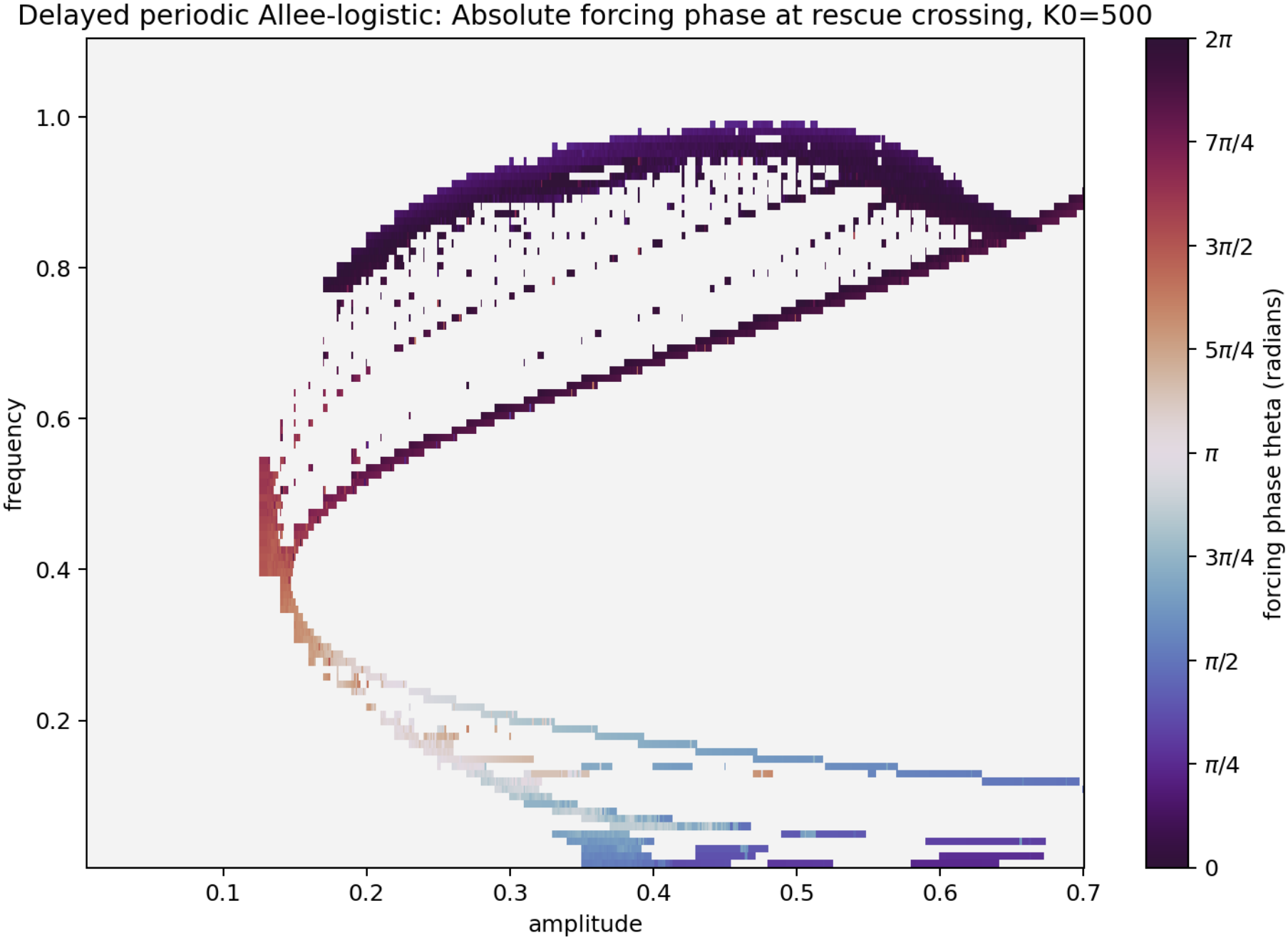
Circular mean of the carrying-capacity phase at successful rescue initiation for K0 = 500.

See Table 2 for the parameter-bank definitions.

**Table 2.** Parameter banks used to isolate Allee-threshold, abundance-scale, and delayed-feedback effects. F, K, and M refer to the respective experiment series.

| **Bank** | *K*<sub>0</sub> | *A* | *r*<sub>0</sub> | **Initial population**<br>*N*<sub>0</sub> | *R*=*K*<sub>0</sub>/*A* |
|---|---|---|---|---|---|
| F01 | 500 | 166.666667 | 1 | 300 | 3.00 |
| F02 | 500 | 131.578947 | 1 | 300 | 3.80 |
| F03 | 500 | 123.456790 | 1 | 300 | 4.05 |
| F04 | 500 | 119.047619 | 1 | 300 | 4.20 |
| F05 | 500 | 113.636364 | 1 | 300 | 4.40 |
| F06 | 500 | 100 | 1 | 300 | 5.00 |
| F07 | 500 | 90.909091 | 1 | 300 | 5.50 |
| K02 | 380 | 100 | 1 | 228 | 3.80 |
| K04 | 420 | 100 | 1 | 252 | 4.20 |
| K07 | 550 | 100 | 1 | 330 | 5.50 |
| M_alpha_0p15 | 500 | 75 | 0.705882353 | 300 | 6.666667 |
| M_alpha_0p25 | 500 | 125 | 1.333333333 | 300 | 4.00 |
| M_alpha_0p3 | 500 | 150 | 1.714285714 | 300 | 3.333333 |

#### 4.2.1 Varying A across Hopf bifurcation thershold

Below the Hopf threshold (F01–F03), surviving population responses were predominantly synchronized with the external forcing, with limited quasiperiodic organization and no well-developed hierarchy of resonance tongues. Preliminary Floquet-multiplier analysis nevertheless indicated a weakening of lyapunov stability in regions preceding the emergence of these structures(figure 9), in other words,even when the system is not yet internally oscillatory, there is a clear resonance behavior(critical slowing down) leading to extinction. Immediately beyond the Hopf threshold (F04, (R=4.20)), the parameter space underwent a qualitative reorganization, with the emergence of extensive low-Lyapunov nonperiodic regions interspersed with distinct periodic-locking tongues. This organization became increasingly pronounced as (R) increased further (F05– F07), consistent with the onset of autonomous oscillations enabling interaction between intrinsic and external frequencies.

**Figure 9.**
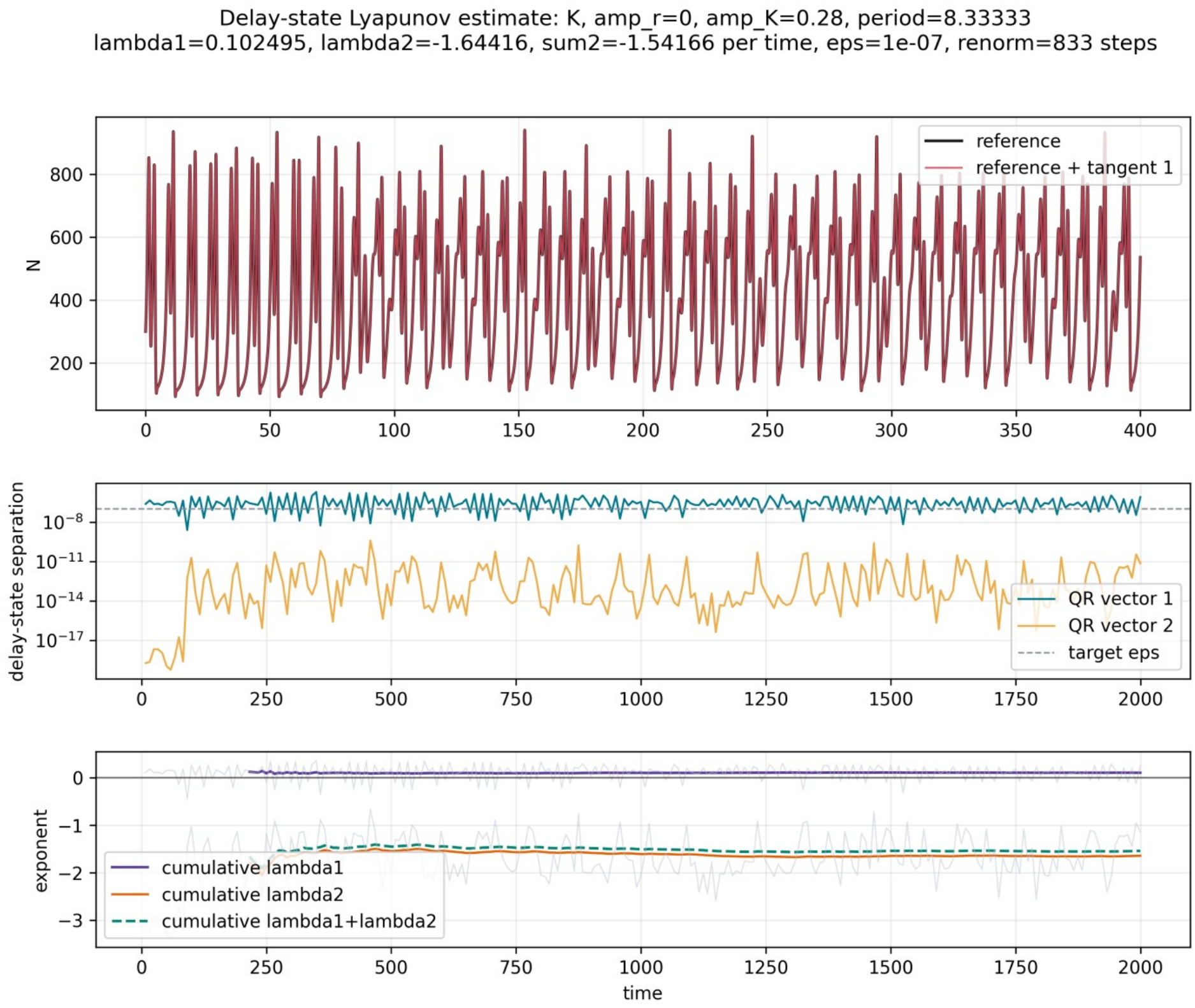
example of a single run’s lyapunov trajectory calculation

**Fig. 10.**
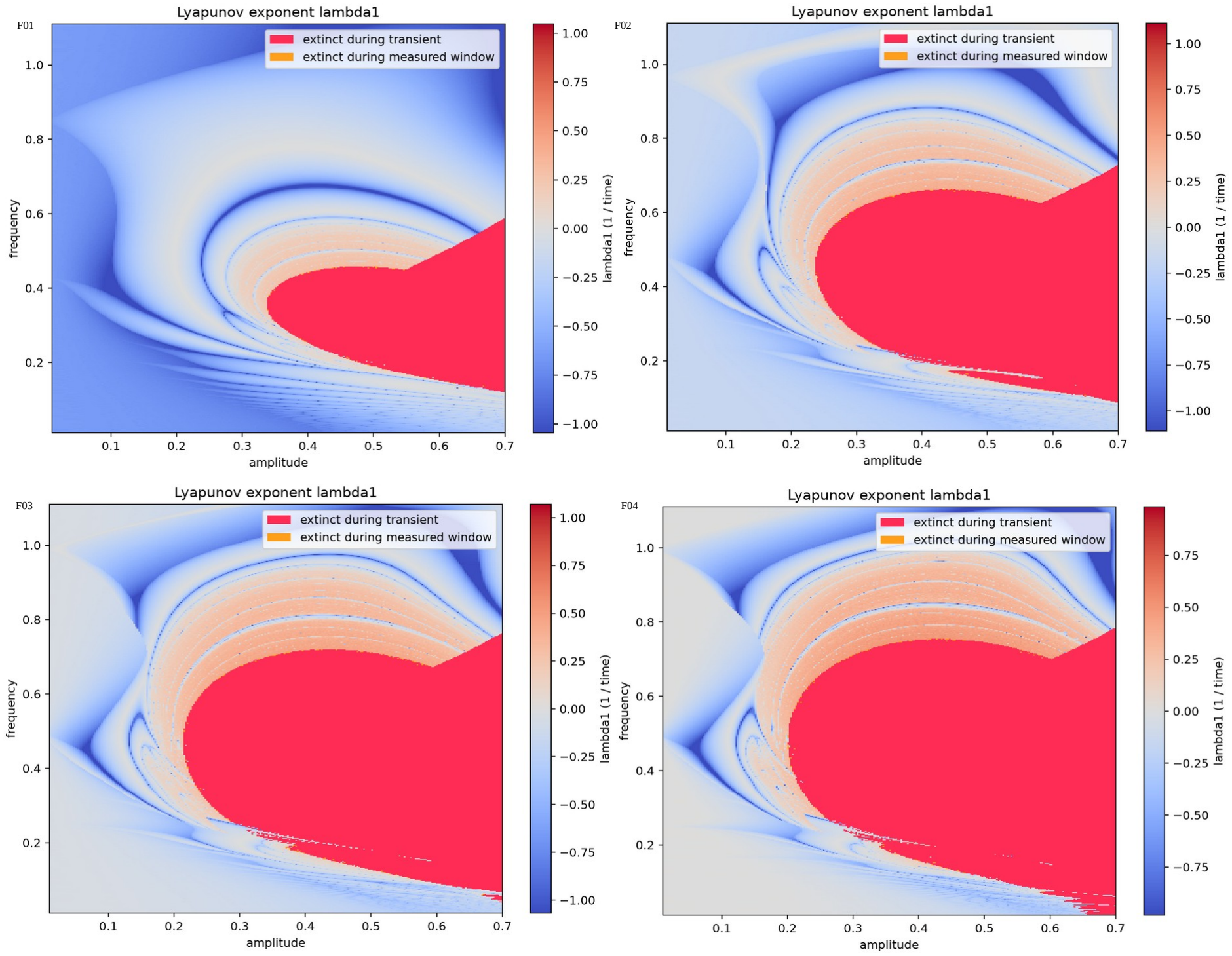
Map of the leading finite-time Lyapunov exponent over forcing amplitude and frequency in the pre-Hopf-delay experiment. Extinction is distinguished by the pink/orange masks; nonextinct cells are colored by the exponent value. Parameter values for the pre-Hopf delay should be specified before submission.

**Fig. 11.**
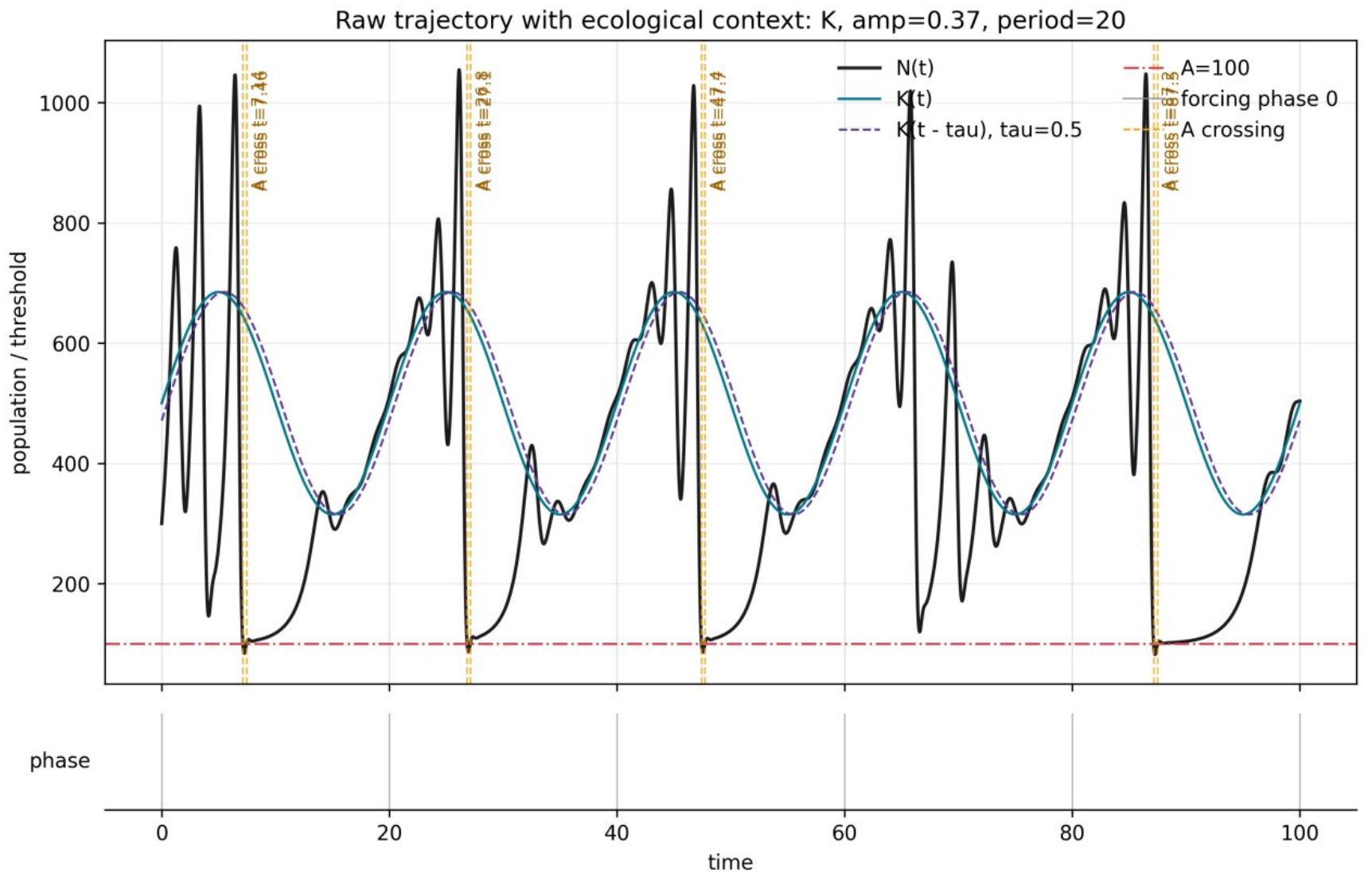
Illustrative population trajectory with seasonal carrying-capacity forcing, delayed abundance, Allee threshold and marked downward threshold crossings. The curve illustrates repeated dives and possible recovery in a single simulated realization (the plotted settings report K amplitude ≈ 0.37 and period = 20).

In contrast to the threshold-like emergence of resonance organization, the extinction boundary shifted progressively toward smaller forcing amplitudes across F01–F07, including before the Hopf threshold was crossed. The broad extinction domain remained concentrated around the principal oscillatory frequency region. Notably, decreasing the Allee threshold was accompanied by an expansion of the extinction domain, despite reducing the critical population density associated with the strong Allee effect. These results distinguish the direct demographic role of the Allee threshold from its influence on the underlying population dynamics, suggesting that increased dynamical instability can outweigh the direct persistence advantage of a lower threshold.

#### 4.2.2 The carring-capcaity-to-A ratio and hopf bifurcation component

Varying (K_0) while preserving (R=K_0/A) produced dynamical maps closely resembling their corresponding F-series cases (K02–F02, K04–F04, and K07–F07). This agreement is consistent with the scale invariance predicted by the nondimensionalized model.

compensatory variation of (r_0) and (A), approximately preserving the linearized delayed-feedback coefficient (c=r_0(R-1)), reproduced much of the resonance and extinction structure of F06 across the M-series. Minor differences remained in the weak-forcing regions and the boundaries of irregular dynamical responses.

Together, these comparisons indicate that the global dynamical organization is strongly associated with the composite parameters governing local delayed-feedback stability. We see in F01-F07 that the frozen hopf bifurcation component clearly divide parameter space behaviour even in high-frequency forcing regions, and a restoration in K and r clearly reverses the effect, highlighting the role of A that participating in exctinction boundary determination through hopf bifurcation threshold against its trivial effect of raising extinction boundary.

### 4.3 Phase-dependent recovery after crossing below the nominal Allee threshold

To investigate how delayed population regulation and periodic environmental forcing influence extinction, we examined population trajectories near the extinction boundary. We identified a recovery mechanism through which populations can temporarily fall below the nominal Allee threshold without necessarily proceeding toward extinction. This occurs because population growth depends on past rather than current abundance, creating a time interval during which the population can recover despite having crossed below the threshold. Figure 6 illustrates the sequence of population decline and possible recovery.

Initially, the population tracks the periodically varying carrying capacity around a stable equilibrium (Figure 6, stage 1). As carrying capacity increases, the equilibrium may lose stability through a Hopf bifurcation, producing larger population oscillations (stage 2). When carrying capacity subsequently declines, the population cannot immediately adjust, and its delayed abundance may remain above the instantaneous carrying capacity. According to the per-capita growth rate q(t) defined in Eq. (11), N(t − *τ*) > K(t) produces negative growth, driving the population downward and potentially across the Allee threshold at t1. However, as carrying capacity rises relative to the delayed abundance, the condition A0 < N(t − *τ*) < K(t) can be restored, allowing q(t) to become positive even while the current population remains below A. The population consequently reaches a minimum at t2 before beginning to recover (Figure 6, stage 3).

The success of this recovery depends on whether the accumulated positive growth is sufficient to compensate for the preceding decline. As demonstrated by the integrated log-growth relationship in Eq. (12), the change in population abundance is determined by the cumulative per-capita growth rate rather than its instantaneous sign. In the successful recovery trajectory, positive growth restores abundance above A, reaching a local maximum at t3. However, the earlier period of low abundance can subsequently enter the delayed feedback term, producing another interval of negative growth and a secondary minimum at t4. If population abundance remains above A through this secondary decline, recovery can continue. Conversely, if negative growth persists and the population fails to recover before delayed low abundance reinforces its decline, the trajectory may proceed toward extinction (Figure 6, stage 4). This illustrates how delayed feedback creates both an opportunity for recovery and a subsequent risk of renewed decline following an initial threshold crossing.

We further examined whether successful rescue events occur at particular phases of environmental forcing. Figure 7 shows the circular mean of the carrying-capacity phase at the initial downward threshold crossing for successful recovery events, calculated according to Eqs. (13)–(14). The resulting map exhibits coherent bands across the forcing-amplitude–frequency parameter space, indicating that the characteristic phase associated with successful recovery varies systematically with the forcing frequency and amplitude, where higher frequency corresponds to later in the phase, and higher amplitude correspond to either start or end of the phase. Within individual parameter combinations, successful recovery events are generally concentrated around similar phases, as indicated by their low circular variance.

Together, these results suggest that the timing of population decline relative to the environmental forcing cycle is an important feature of the recovery process. The delayed response creates a temporary opportunity for recovery following a threshold crossing, while the phase statistics characterize how the timing of successful recovery varies across environmental forcing conditions.

## 5. Discussion

Population persistence under environmental variability depends not only on the intensity of external disturbances but also on how those disturbances interact with internal population dynamics. Our results illustrate three interconnected ecological principles: resonance between environmental and endogenous oscillations can determine extinction vulnerability; delayed density dependence makes population recovery dependent on recent history and environmental timing; and conditions that increase potential population growth can also destabilize populations. Together, these findings connect the dynamical organization of periodically forced populations to broader questions of ecological resilience and extinction.

### 5.1 Ecological resonance, how it promotes extinction and disconnect regularity and persistence

The interaction between endogenous population cycles and periodic environmental variability has long been recognized as an important determinant of population dynamics..Our findings extend this framework by demonstrating that resonance can organize not only the temporal behavior of a population but also its vulnerability to extinction. In the investigated parameter space, environmental variation was particularly damaging when its frequency approached the population’s principal endogenous oscillation frequency. Extinction could occur even when the minimum environmental carrying capacity remained above the Allee threshold, indicating that environmental deterioration alone was insufficient to explain collapse. Instead, resonance amplified population fluctuations sufficiently for their interaction with low-density feedback to produce extinction.

An important implication is that the ecological consequences of environmental variation depend on its timing relative to intrinsic population dynamics, rather than simply on its amplitude. Environmental fluctuations of similar magnitude can therefore have substantially different consequences for populations with different endogenous rhythms. Moreover, changing the strength of seasonal variation, even without changing its frequency, may alter whether a population becomes synchronized with its environment and consequently change its extinction vulnerability.

Luo et al. (2026), demonstrated that frequency locking can suppress destructive oscillations and promote coexistence in phage–bacteria systems. Our results provide a complementary example in which resonance can increase extinction vulnerability through its interaction with a strong Allee effect. The comparison suggests that synchronization itself is neither inherently stabilizing nor destabilizing; its ecological consequences depend on the population’s underlying regulatory mechanisms and the extent to which synchronized fluctuations approach extinction thresholds.

This distinction is also important when interpreting complex population dynamics. Our tongue dominated parameter space simulations show that greater irregularity region does not necessarily correspond to greater extinction vulnerability, it might sometimes indicate exact opposite.

In a resonance tongue organized parameter space, periodic-locked region become wider as amplitude grows and low amplitude regions are generally more quasiperiodic. A population exhibiting relatively regular oscillations may be more susceptible to collapse than one with more complicated dynamics. In the high-frequency region, we see chaostic regions expand as extinction boundary retreats, in lower extinction region, we see smaller and smaller tongues only surrounding the extinction boundary. Consequently, temporal regularity alone may be a poor indicator of ecological persistence. Understanding environmental vulnerability requires considering both the organization of population fluctuations and the mechanisms through which those fluctuations lead to population decline.

### 5.2 Allee effects and ecological resilience: the importance of history and environmental timing

The Allee effect introduces positive density dependence at low population abundance, creating conditions under which population decline can become self-reinforcing. This mechanism is commonly associated with critical population thresholds separating persistence from extinction. However, we demonstrated that in our delayed-feedback systems population can be rescued after crossing threshold, and the rescue events concentrates at a phase determined by forcing frequency and amplitude.

Previous studies have demonstrated that the interaction between delayed regulation and Allee effects can produce extinction, bistability, and sensitivity to initial conditions (Liz and Ruiz-Herrera 2015). Our results extend this understanding by demonstrating that temporary declines below the nominal Allee threshold need not result in irreversible population collapse. Because the growth response depends on past population abundance, a population may retain opportunities for positive growth even after its current abundance has fallen below the threshold.

The fact that successful recovery events were frequently concentrated within restricted phases of the environmental cycle, suggests that ecological resilience depends not only on how far a population declines, but also on when that decline occurs relative to environmental variation. Two declines reaching similarly low population abundance may have different consequences if they occur under different environmental timing, because the subsequent opportunities for recovery can differ.

The importance of timing has also emerged in recent studies of ecological tipping points. Alkhayuon et al. (2021) demonstrated that transitions from cyclic population dynamics to extinction can depend on the phase of a population cycle when environmental conditions change. Although their phase-tipping framework concerns a different form of environmental transition, both studies illustrate the broader principle that ecological outcomes may depend on the temporal relationship between internal dynamics and external environmental change. Our findings extend this perspective by identifying environmental-phase-associated recovery opportunities during recurrent forcing.

These results also suggest a distinction between temporary threshold crossing and long-term extinction. A decline below a nominal critical abundance is not necessarily sufficient to determine population fate when delayed feedback remains important. Instead, persistence depends on whether favorable conditions occur early enough, and for sufficiently long, to restore population growth.

From an ecological perspective, this suggests that assessments of population vulnerability based solely on current abundance may overlook important information. Recent population history and environmental timing may also be relevant to evaluating recovery potential. Nevertheless, the observed concentration of successful rescue events does not yet establish a universal favorable phase or an effective intervention window. Identifying such windows would require further investigation of how recovery responds to changes in environmental timing and population history.

### 5.3 The paradox of enrichment: how favorable conditions can destabilize populations

A longstanding problem in theoretical ecology is that environmental conditions favorable to population growth do not necessarily promote population stability. The classical paradox of enrichment describes how increasing resource availability can destabilize consumer–resource systems, potentially producing large fluctuations that threaten persistence (Rosenzweig 1971). Related counterintuitive outcomes have also been identified in single-population models with Allee effects, where increasing carrying capacity or mating success can sometimes promote extinction rather than persistence (Schreiber 2003).

Our results connect this broader ecological principle to the interaction between delayed density dependence, Allee effects, and environmental forcing. In the present model, the ratio between carrying capacity and the Allee threshold influences not only the range of population abundances compatible with positive growth but also the strength of the feedback regulating population abundance. Changes in this ratio can therefore modify the stability of the population’s equilibrium and determine whether endogenous oscillations emerge.

This creates a mechanism through which conditions associated with greater potential population growth can also increase dynamical vulnerability. Stronger delayed feedback can produce larger endogenous fluctuations, which subsequently interact with periodic environmental forcing. Once intrinsic oscillations become established, resonance provides an additional pathway through which environmental variability can generate sufficiently large declines to threaten persistence.

The results therefore illustrate an enrichment-related destabilization principle without requiring an explicitly modeled predator–prey interaction. Unlike the classical paradox of enrichment, where destabilization arises through trophic interactions, the mechanism considered here operates through delayed regulation within a single population. The Allee effect further connects this destabilization to extinction by making sufficiently severe population declines difficult to reverse.

An additional consequence is that lowering the Allee parameter does not necessarily improve persistence in this particular formulation. Although a lower threshold might be expected to reduce extinction vulnerability, it also changes the strength of density-dependent growth and can enhance destabilizing feedback. Thus, the ecological consequences of modifying an Allee parameter depend on whether that modification affects only the critical density or also alters population regulation.

Our parameter comparisons support the importance of this coupled effect, although the direct consequences of increasing carrying capacity independently of the Allee parameter require further testing. More broadly, these findings emphasize that favorable environmental conditions and demographic stability are not equivalent. Changes that increase potential equilibrium abundance may also strengthen endogenous fluctuations and modify their resonance with environmental variability, producing unexpected consequences for population persistence.

### 5.4 Broader implications and limitations

Together, these three mechanisms suggest that extinction vulnerability emerges from interactions among environmental variation, endogenous population regulation, and low-density feedback. Resonance determines which environmental frequencies and amplitudes produce particularly strong population responses; delayed density dependence influences the timing and possibility of recovery; and the strength of demographic feedback determines whether those responses remain stable or become strongly oscillatory.

These conclusions arise from a deterministic, single-population model with fixed initial history and idealized periodic forcing. Natural populations experience stochastic disturbances, multiple interacting regulatory mechanisms, and environmental variability that is rarely perfectly periodic. Moreover, some dynamical transitions may depend on initial population histories or coexisting attractors, which require further analysis to distinguish. The present results therefore identify mechanisms of persistence and extinction rather than universal predictions for particular ecological systems.

Nevertheless, the study highlights a general ecological implication: predicting population responses to environmental change requires attention not only to the magnitude of external disturbances or the location of static population thresholds, but also to the temporal relationship between environmental forcing, internal population rhythms, and opportunities for recovery.

## Acknowledgments

Junjie Sun acknowledges summer research funding from the Centre for Applied Mathematics in Bioscience and Medicine (CAMBAM), McGill University.

## Code availability

Related code repository: https://github.com/jsun76/intrinsic-vs-external-forcing-hutchinson-model

## Appendix A. Autonomous forcing clock

For periodic-orbit calculations, the explicit time dependence can be represented by a two-dimensional attracting clock:

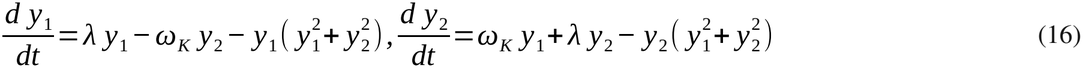

In polar coordinates, 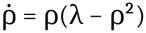and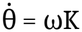. With λ = 1 the clock approaches ρ = 1. The implemented lifted model uses φK = −π/2 and K = K0 + BKy2 on the clock cycle, which is equivalent to equation (2). The population component remains delayed; the clock only converts the forcing into an autonomous subsystem.This is used in fold curve extension in DDE-Biftool, as the software requires autonomous equations. (Sieber et al. 2017).

## Appendix B. Numerical implementation and diagnostic criteria

### B.1 Raw run labeling and extinction condition

In each run we monitor several independent attributes, after detecting one of which is run will be cut off (except repetitive Allee crossings) (Table 3).

**Table 3.** Attributes retained for early-stopping rules, numerical extinction, and Allee-threshold crossing events.

| **Attribute** | **Implemented meaning** |
|---|---|
| stop_reason | Requested step limit reached, numerical zero reached, or online periodic hit |
| periodic_hit | A recurrence was accepted; order, method, errors and matched times are recorded |
| effective_extinct | Population reached the positive extinction cutoff |
| literal_extinct | Population reached zero or below |
| Allee crossing/rescue fields | Downward/upward crossings of *A*, paired rescue events and unresolved excursions |

Effective extinction uses first passage across the cutoff in the **entire computed record, including the transient**:

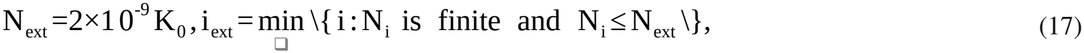

Downstream analysis distinguishes transient extinction from measurement-period extinction using whether this index precedes the transient boundary.

#### B. History-state periodicity detection

The detector identifies candidate periodic responses by comparing the **complete sampled population history over one delay interval**, rather than comparing individual population values.

The mathematical criterion is:

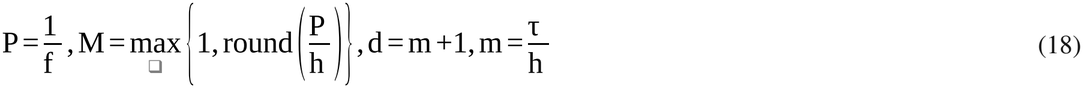

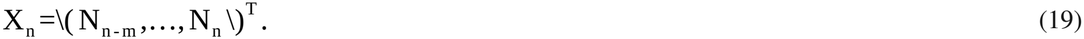

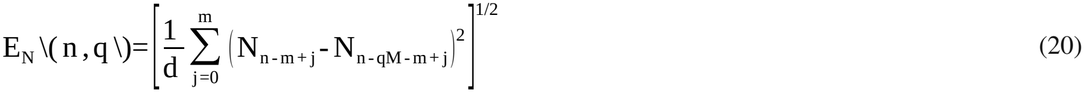

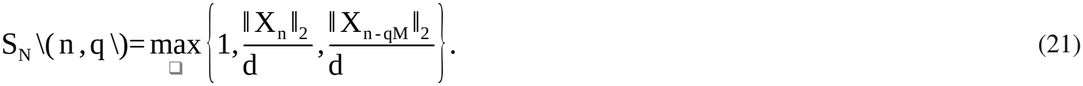

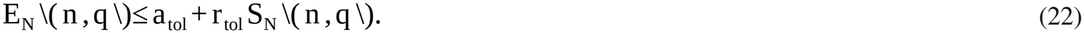

Here, q is the number of forcing cycles separating the compared windows. The online detector checks once per rounded forcing period after the transient, testing orders 1-8.

Most of the heatmpa settings use:

- Relative tolerance: 1e-5.
- Absolute tolerance: 2e-9 × K0
- Minimum check endpoints: two.

The first accepted online comparison stops the raw integration. Afterwards, another search compares histories separated by exact forcing periods, using linear interpolation of the fixed-step solution. A successful interpolated result replaces the online result; if interpolation finds no hit, an existing online hit is retained. The resulting label therefore means observed numerical recurrence within the specified tolerances.

#### B.3 Lyapunov-exponent implementation

The reference population obeys:

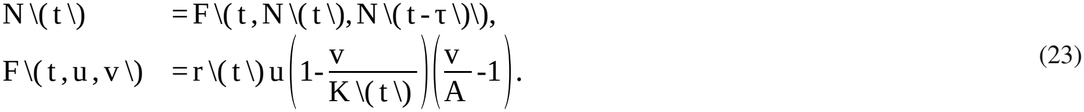

An infinitesimal perturbation z(t) satisfies the linear variational equation:

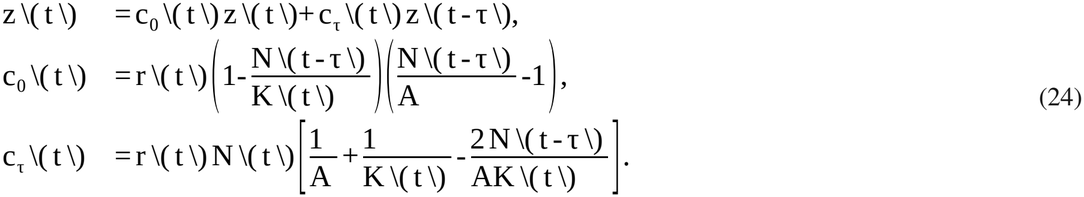

These coefficients are obtained by differentiating the model with respect to its current and delayed population arguments. The external forcing schedule remains identical for all perturbations.

Because the model contains delay, its numerical state includes the whole sampled history window:

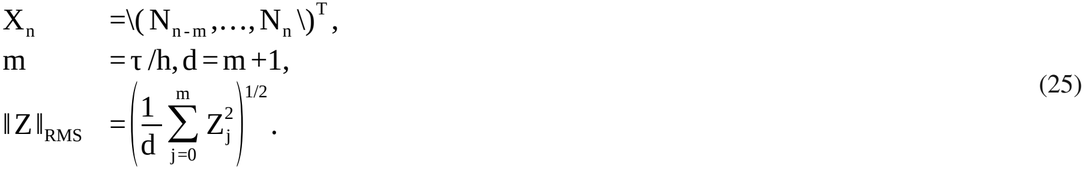

For the default parameters, h=0.01 and tau=0.5, this state has **51 coordinates**. Two independent tangent histories are propagated within that state.

The reference trajectory and tangent histories are advanced together using the same fixed-step Heun scheme. Tangent coefficients are evaluated at the corresponding predictor and corrector stages. At each orthogonalization endpoint, the tangent histories form a matrix whose columns contain their complete delay windows:

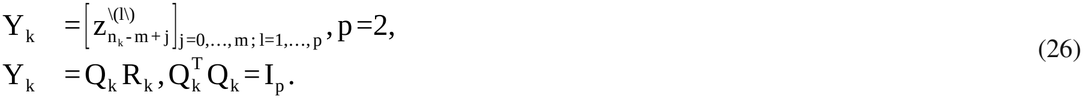

Each tangent column is then restored to its prescribed RMS magnitude. With a positive-diagonal QR convention:

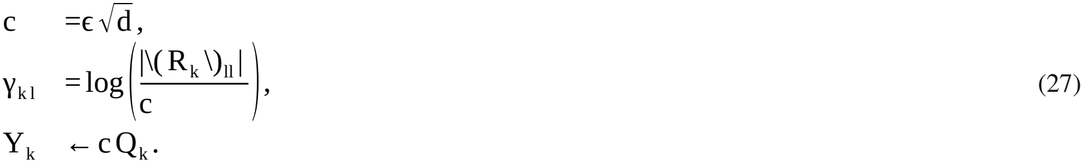

The implementation uses absolute diagonal values and an equivalent sign correction. **The reset replaces the entire current perturbation-history window**, rather than only its present value.

The finite-horizon exponent estimate is:

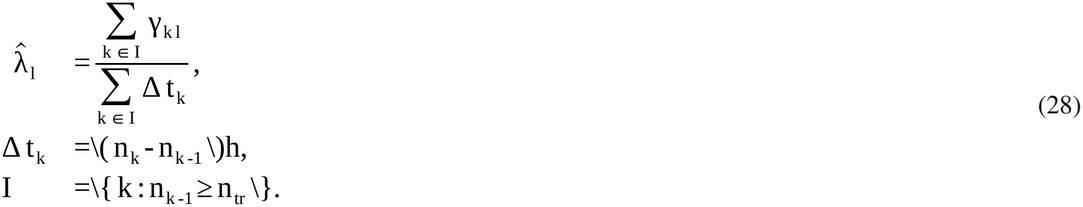

Orthogonalization continues during the transient, but only blocks starting after the transient contribute to the average. The denominator is the **included averaging time**, rather than the requested total simulation duration. The discrete QR construction and logarithmic accumulation are described explicitly in Breda and Liessi, Section 2, Eqs. (2.3)–(2. (Breda and Liessi 2024).

### B.4 Floquet multiplier computation

For a candidate periodic response, Floquet multipliers measure how perturbations change over one complete response period. For an order-*q* response, this corresponds approximately to *q* forcing periods. (Breda et al. 2006).

The theoretical construction uses the evolution of perturbations in delay-history space over one period, called the monodromy operator. Its eigenvalues are the Floquet multipliers. This framework is described by Breda, Maset and Vermiglio (2006). Our implementation constructs a finite matrix using the existing fixed-step Heun discretization. (Breda et al. 2006).

For the delayed population equation, the sampled history and linear variational equation are

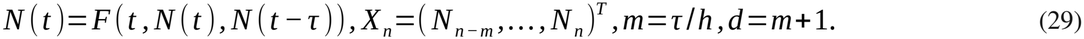

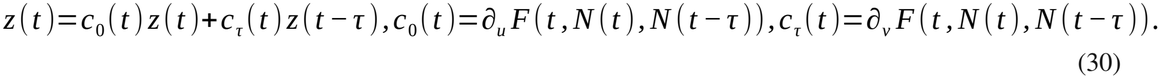

The coefficients are evaluated analytically at both Heun stages. Each perturbation follows the same discretization as the reference trajectory:

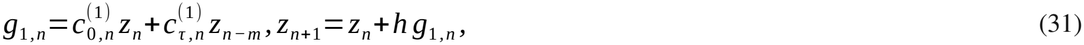

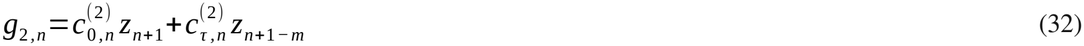

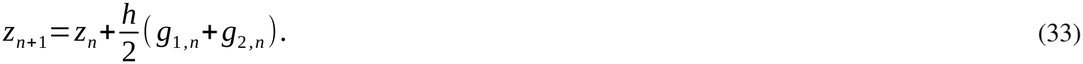

The entire perturbation history shifts forward after each update.

Unlike the two-column Lyapunov calculation, Floquet analysis propagates **every basis direction** of the sampled history state. Starting from the identity matrix, it constructs the derivative of the complete return map:

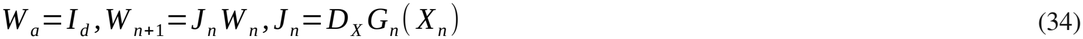

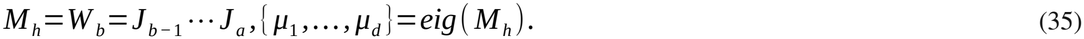

Here, *G*_*n*_ is the Heun history update and *a, b* delimit the selected response interval. No QR reset is applied during this propagation.

With the current dt=0.01 and delay 0.5, the matrix is **51 × 51**, producing 51 complex multipliers. These constitute the full spectrum of the discrete history map.

#### B.5 Stability interpretation and current labels

For a genuinely periodic response, a mode decays when its multiplier lies inside the unit circle and grows when it lies outside. Autonomous nonconstant cycles have a neutral time-shift multiplier at +1, which is excluded when assessing orbital stability. This interpretation is documented in DDE-BIFTOOL’s manual, Section 5.1.2. The current F/K/M experiments have externally imposed forcing with fixed phase, so the calculation does not introduce a neutral forcing-clock direction.

The code uses the largest relevant multiplier modulus:

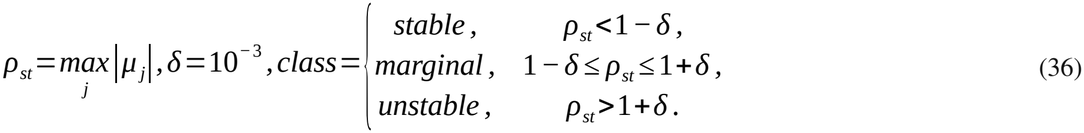

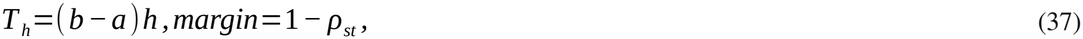

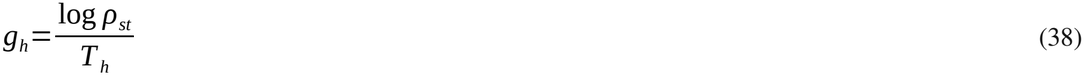

The growth rate permits comparison across different response periods. It is calculated from the one-period matrix, whereas the Lyapunov estimate accumulates growth over a longer trajectory.

All multipliers remain stored. The 1e-8 significance cutoff only controls how many are counted as significant.

#### B.6 MATLAB continuation protocol

Periodic solutions and their folds are computed using DDE-BIFTOOL 3.1.1. (Engelborghs et al. 2002; Sieber et al. 2017)).

The periodically forced delayed Allee model is

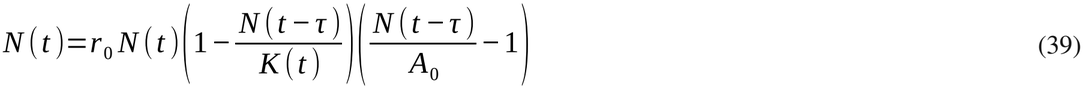

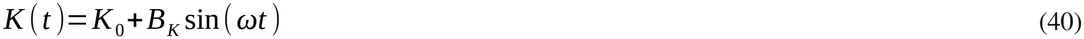

The parameter-space coordinates and response period are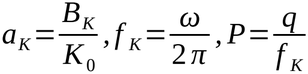, where *q* is the smallest positive integer for which the complete delayed history repeats. Initial periodic profiles are obtained by direct simulation and subsequently corrected as periodic boundary-value problems.

The forcing is embedded in an autonomous system by introducing a two-dimensional clock. Denoting the lifted state by (*x*=(*N, y*_1_, *y*_2_)^*T*^), the system is written as 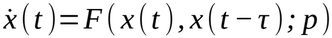.For a periodic orbit (*x*_∗_(*t*)) of period *P*, the normalized profile (*u* (*s*)=*x*_∗_(*Ps*))satisfies

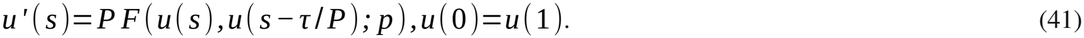

Delayed arguments are evaluated using periodic extension. An integral phase condition removes the arbitrary temporal shift. The orbit is represented by piecewise degree-four polynomials and corrected by Newton iteration using Gauss--Legendre collocation points.

Floquet multipliers are calculated from the variational equation along the corrected orbit:

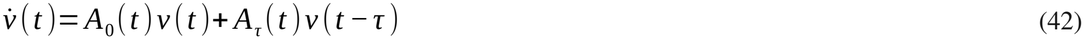

where

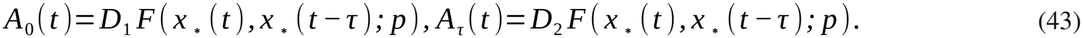

For population perturbations with forcing phase held fixed, the corresponding scalar coefficients are

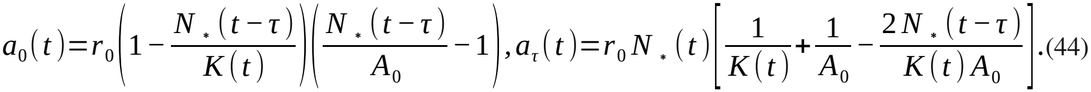

The implementation evaluates the full lifted variational system, including clock perturbations.

Let (*ϕ*) denote an initial perturbation history on ([−*τ*, 0]). The monodromy operator maps this history through one response period:

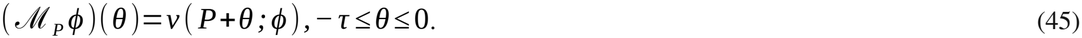

Its eigenvalues are the Floquet multipliers:

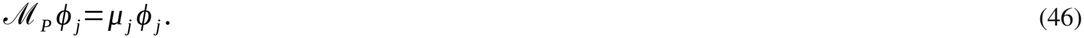

DDE-BIFTOOL approximates this history-to-history operator by discretizing the linearized collocation equations and calculating the eigenvalues of the resulting finite-dimensional map.

The corresponding real Floquet growth rates are

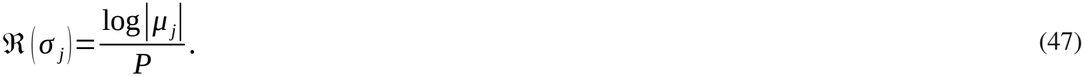

The autonomous formulation contains a trivial multiplier at +1 associated with temporal phase. This multiplier is excluded when assessing orbital stability. A generic fold of periodic orbits contains an additional multiplier at +1:

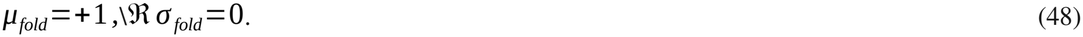

Consequently, a zero real growth rate alone is insufficient to identify a fold: period doubling involves a multiplier at − 1, whereas a torus bifurcation involves a nonreal conjugate pair on the unit circle. Fold candidates are identified from nontrivial multipliers approaching +1, together with a turning point along a one-parameter periodic-orbit branch.

Candidate folds are corrected using the SetupPOfold extended system. Schematically, the fold-defining equations are

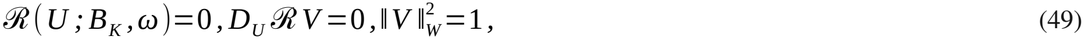

where *R* contains the periodic-orbit equations and phase condition, *U* includes the orbit and period, and *V* represents the additional null direction.

Allowing both *B*_*k*_ and *ω* to vary defines a one-dimensional fold curve.

Both ends of each local fold branch are extended using secant prediction and Newton correction with a pseudo-arclength condition. Writing the extended unknowns and free parameters as *Z*, the continuation step is represented by

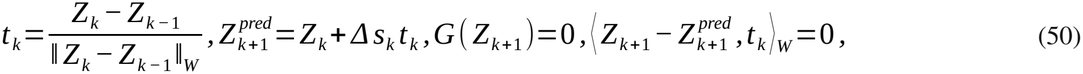

where *G*=0 denotes the fold-defining extended system.

The stepsize decreases after difficult or failed corrections and increases after consistently successful continuation. Adaptive remeshing and backtracking provide recovery attempts.

For the low-frequency preset, maximum parameter increments are

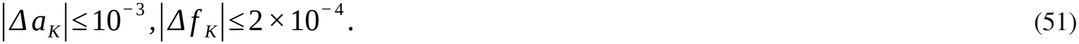

Initial meshes contain 160 intervals, increasing to 240 for response periods exceeding 15 time units; adaptive refinement permits up to 720 intervals. Each direction has a budget of 2400 continuation points and 120 additional recovery points.

Every global fold point is audited by extracting the original periodic orbit and recomputing its equation defect and Floquet multipliers. The configured acceptance thresholds are

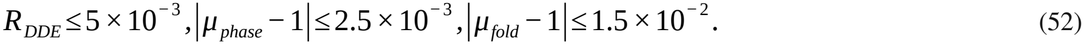

where *RDDE* is the maximum sampled equation residual, scaled componentwise by 1+|*F*_*i*_|. Remaining-spectrum stability is assessed after removing both the phase and target fold multipliers.

Continuation stops at a configured domain boundary, a numerical return consistent in parameters, period and orbit profile, or an unresolved numerical stopping condition. Stagnation and exhausted budgets are recorded explicitly. Untrusted points are not joined in the plotted curves, and numerical termination is not treated as evidence that a resonance tongue has been completely enclosed.

## Appendix C. Supporting figures

### C.1 Examples of Run classification

**Figure 10.1.**
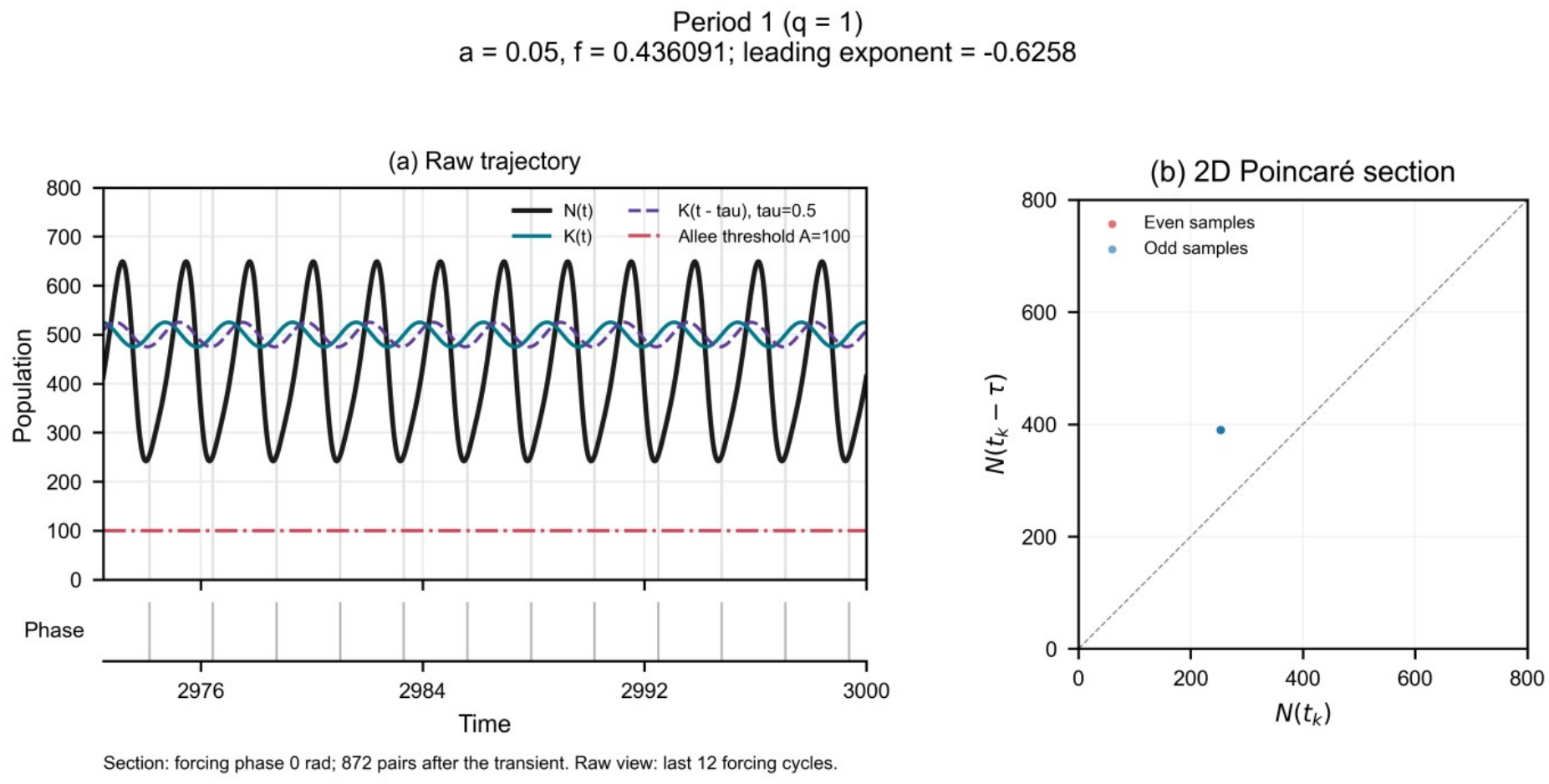
Run classified as q=1 periodic

**Figure 10.2.**
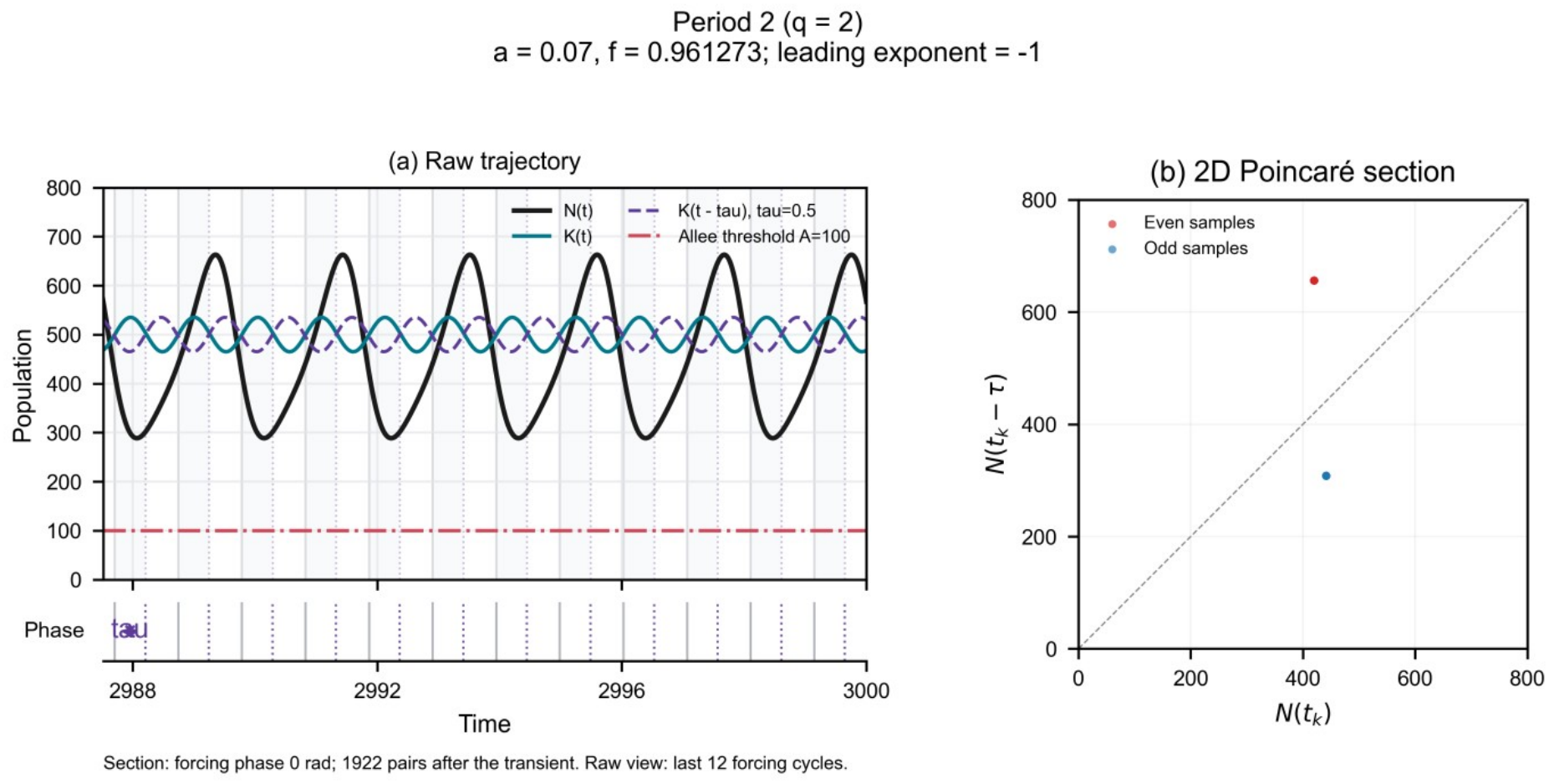
Run classified as q=2 periodic

**Figure 10.3.**
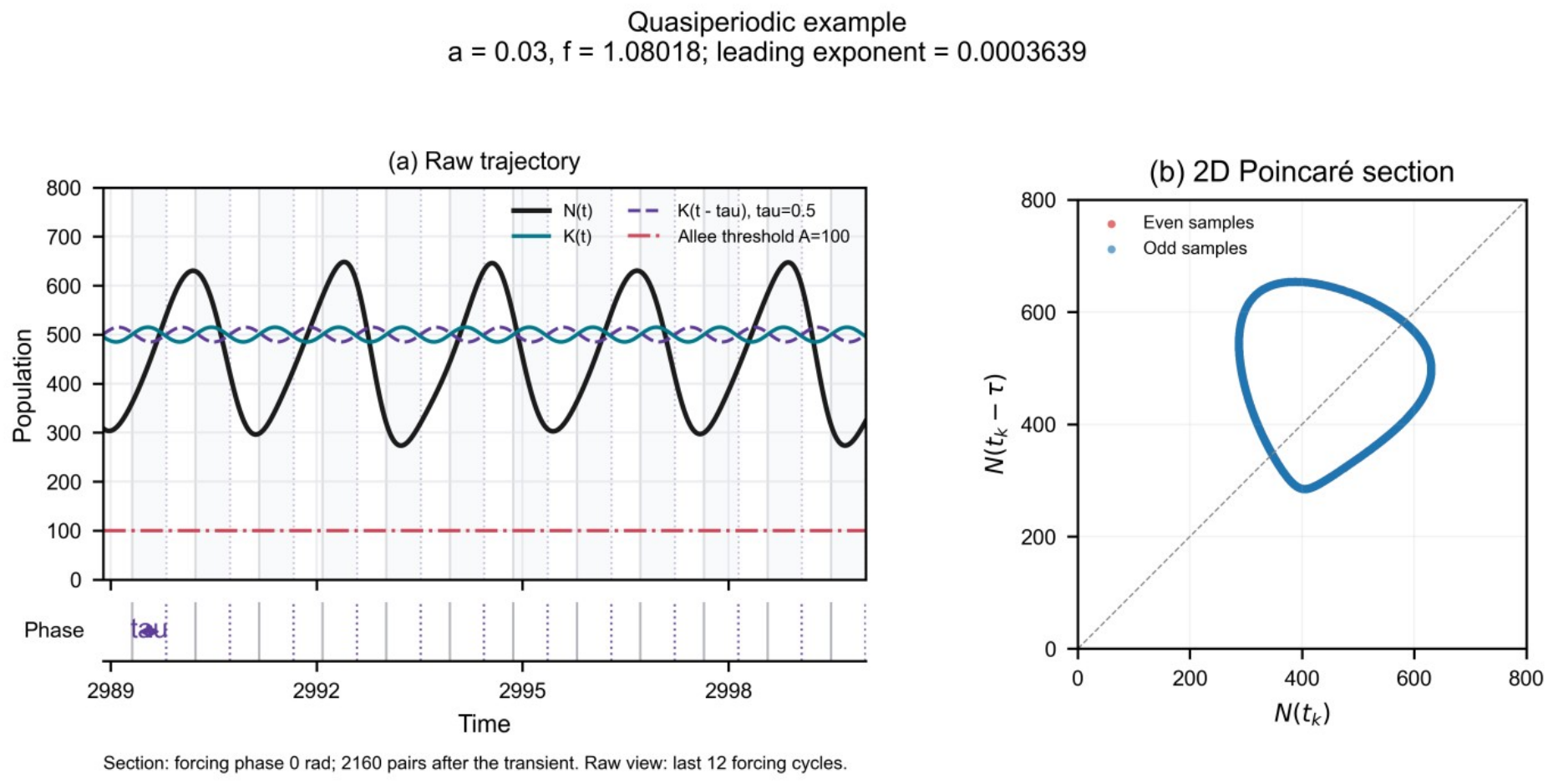
Run classified as quasiperiodic

**Figure 10.4.**
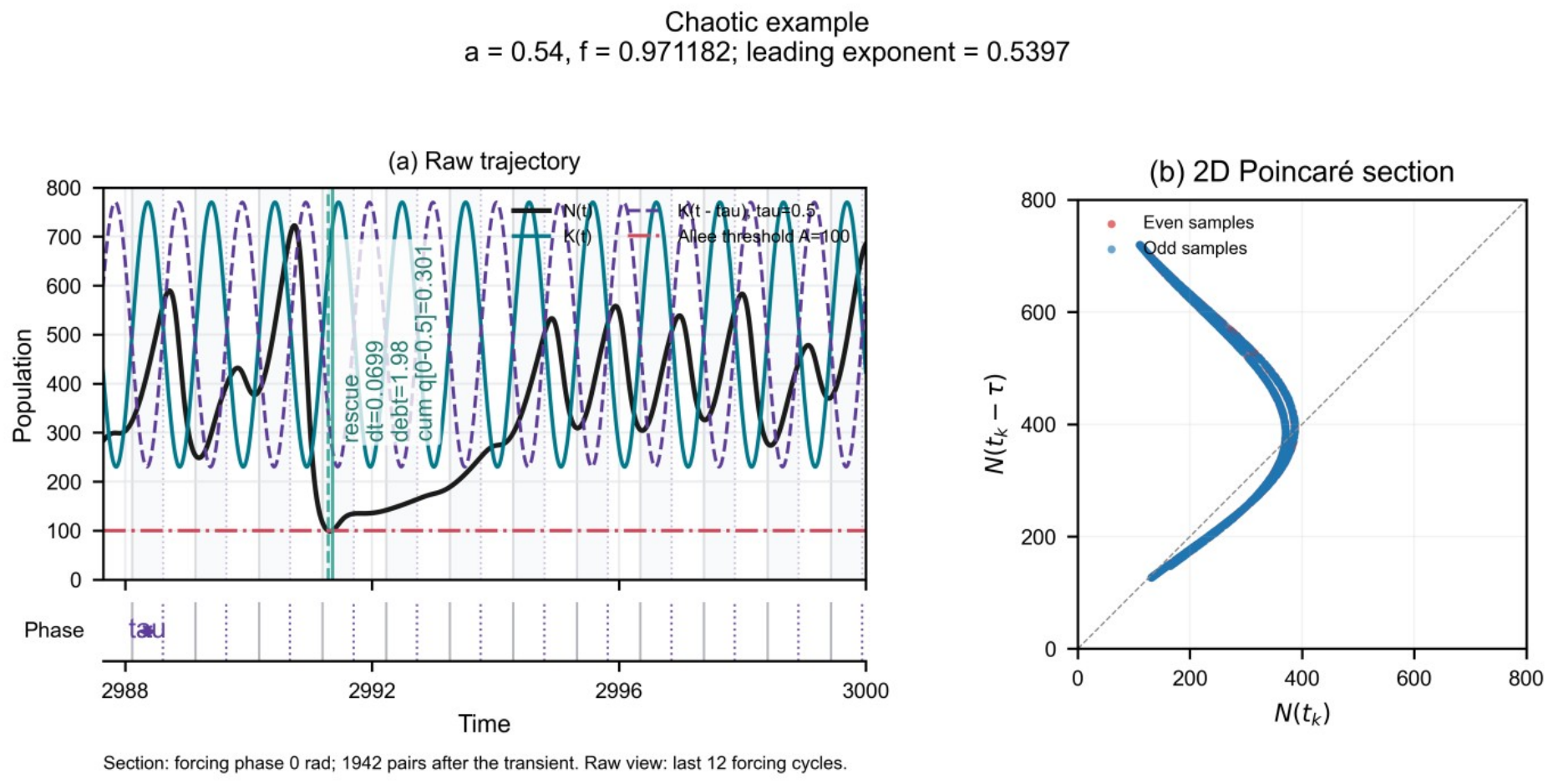
Run classified as chaotic with a rescue event

### C.2 Critical Slowing down in F01-F03, Pre Hopf bifurcation

### C.3 Illustrative rescue trajectory

An example trajectory showing successive Allee crossings and recoveries is provided in Fig. 10.

## References

Alkhayuon H, Tyson RC, Wieczorek S (2021) Phase tipping: how cyclic ecosystems respond to contemporary climate. Proc R Soc Math Phys Eng Sci 477:20210059. 10.1098/rspa.2021.0059

Allen JC (1990) Chaos and phase-locking in predator-prey models in relation to the functional response. Fla Entomol 73:100–110. 10.2307/3495332

Breda D, Liessi D (2024) A practical approach to computing Lyapunov exponents of renewal and delay equations. Math Biosci Eng 21:1249–1269. 10.3934/mbe.2024053

Breda D, Maset S, Vermiglio R (2006) Numerical computation of characteristic multipliers for linear time periodic coefficients delay differential equations

Chugunova MA, Ann Campbell S (2020) Difference population equation with variable Allee effect and periodic carrying capacity. J Differ Equ Appl 26:753–778. 10.1080/10236198.2020.1787396

Courchamp F, Clutton-Brock T, Grenfell B (1999) Inverse density dependence and the Allee effect. Trends Ecol Evol 14:405– 410. 10.1016/S0169-5347(99)01683-3

Engelborghs K, Luzyanina T, Roose D (2002) Numerical bifurcation analysis of delay differential equations using DDE-BIFTOOL. ACM Trans Math Softw 28:1–21. 10.1145/513001.513002

Gurney WSC, Blythe SP, Nisbet RM (1980) Nicholson’s blowflies revisited. Nature 287:17–21. 10.1038/287017a0

Hutchinson GE (1948) Circular causal systems in ecology. Ann N Y Acad Sci 50:221–246. 10.1111/j.1749-6632.1948.tb39854.x

Jankovic M, Petrovskii S (2014) Are time delays always destabilizing? Revisiting the role of time delays and the Allee effect. Theor Ecol 7:335–349. 10.1007/s12080-014-0222-z

Keane A, Krauskopf B, Postlethwaite C (2015) Delayed Feedback Versus Seasonal Forcing: Resonance Phenomena in an El Niñ o Southern Oscillation Model. SIAM J Appl Dyn Syst 14:1229–1257. 10.1137/140998676

Keane A, Krauskopf B, Postlethwaite C (2016) Investigating irregular behavior in a model for the El Niño Southern Oscillation with positive and negative delayed feedback. SIAM J Appl Dyn Syst 15:1656–1689. 10.1137/16M1063605

Liz E, Ruiz-Herrera A, 1. Departamento de Matemática Aplicada II, E.T.S.E. Telecomunicación, Universidade de Vigo, Campus Marcosende, 36310 Vigo, 2. Bolyai Institute, University of Szeged, Aradi vértanúk tere 1., H-6720 Szeged (2015) Delayed population models with Allee effects and exploitation. Math Biosci Eng 12:83–97. 10.3934/mbe.2015.12.83

Luo H-N, Wu Z-X, Guan J-Y (2026) Frequency locking to environmental forcing suppresses oscillatory extinction in phage-bacteria interactions. Phys Rev E 114:014412. 10.1103/rch2-2hsr

Nicholson AJ (1954) An outline of the dynamics of animal populations. Aust J Zool 2:9–65. 10.1071/ZO9540009

Nisbet RM, Gurney WSC (1976) Population dynamics in a periodically varying environment. J Theor Biol 56:459–475. 10.1016/S0022-5193(76)80086-0

Orland MC, Lawler SP (2004) Resonance inflates carrying capacity in protist populations with periodic resource pulses. Ecology 85:150–157. 10.1890/02-0372

Pikovsky A, Rosenblum M, Kurths J (2001) Synchronization: a universal concept in nonlinear sciences. Cambridge University Press

Rosenzweig ML (1971) Paradox of enrichment: destabilization of exploitation ecosystems in ecological time. Science 171:385–387. 10.1126/science.171.3969.385

Schreiber SJ (2003) Allee effects, extinctions, and chaotic transients in simple population models. Theor Popul Biol 64:201– 209. 10.1016/S0040-5809(03)00072-8

Sieber J, Engelborghs K, Luzyanina T, Samaey G, Roose D (2017) DDE-BIFTOOL v. 3.1.1 manual: bifurcation analysis of delay differential equations. DDE-BIFTOOL project

Stephens PA, Sutherland WJ, Freckleton RP (1999) What is the Allee effect? Oikos 87:185–190. 10.2307/3547011

van den Berg JB, Jaquette J (2018) A proof of Wright’s conjecture. J Differ Equ 264:7412–7462. 10.1016/j.jde.2018.02.018

Vandermeer J, Stone L, Blasius B (2001) Categories of chaos and fractal basin boundaries in forced predator–prey models. Chaos Solitons Fractals 12:265–276. 10.1016/S0960-0779(00)00111-9

Vasseur DA, Fox JW (2009) Phase-locking and environmental fluctuations generate synchrony in a predator–prey community. Nature 460:1007–1010. 10.1038/nature08208

Wright EM (1955) A non-linear difference-differential equation. J Reine Angew Math 194:66–87. 10.1515/crll.1955.194.66

